# Estimating encounter location distributions from animal tracking data

**DOI:** 10.1101/2020.08.24.261628

**Authors:** Michael J. Noonan, Ricardo Martinez-Garcia, Grace H. Davis, Margaret C. Crofoot, Roland Kays, Ben T. Hirsch, Damien Caillaud, Eric Payne, Andrew Sih, David L. Sinn, Orr Spiegel, William F. Fagan, Christen H. Fleming, Justin M. Calabrese

## Abstract

1. Ecologists have long been interested in linking individual behavior with higher-level processes. For motile species, this ‘upscaling’ is governed by how well any given movement strategy maximizes encounters with positive factors, and minimizes encounters with negative factors. Despite the importance of encounter events for a broad range of ecological processes, encounter theory has not kept pace with developments in animal tracking or movement modeling. Furthermore, existing work has focused primarily on the relationship between animal movement and encounter *rates* while no theoretical framework exists for directly relating individual movement with the spatial *locations* of encounter events in the environment.
2. Here, we bridge this gap by introducing a new theoretical concept describing the long-term encounter location probabilities for movement within home ranges, termed the conditional distribution of encounters (CDE). We then derive this distribution, as well as confidence intervals, implement its statistical estimator into open source software, and demonstrate the broad ecological relevance of this novel concept.
3. We first use simulated data to show how our estimator provides asymptotically consistent estimates. We then demonstrate the general utility of this method for three simulation-based scenarios that occur routinely in biological systems: i) a population of individuals with home ranges that overlap with neighbors; ii) a pair of individuals with a hard territorial border between their home ranges; and iii) a predator with a large home range that encompassed the home ranges of multiple prey individuals. Using GPS data from white-faced capuchins (*Cebus capucinus*) tracked on Barro Colorado Island, Panama, and sleepy lizards (*Tiliqua rugosa*) tracked in Bundey, South Australia, we then show how the CDE can be used to estimate the locations of territorial borders, identify key resources, quantify the location-specific potential for competition, and/or identify any changes in behaviour that directly result from location-specific encounter probability.
4. This novel target distribution enables researchers to better understand the dynamics of populations of interacting individuals. Notably, the general estimation framework developed in this work builds straightforwardly off of home range estimation and requires no specialised data collection protocols. This method is now openly available via the ctmm R package.

## Introduction

Linking individual behaviour with higher-level processes has long been a cornerstone of ecological research (Darwin, 1859; Skellam, 1951; Hogeweg & Hesper, 1990; DeAngelis & Gross, 1992; Sutherland, 1996; Grimm & Railsback, 2005; Gil *et al.*, 2018). For motile species, this ‘upscaling’ is governed by how well any given movement strategy maximises encounters with positive factors (e.g., food, mates, essential resources), and minimises encounters with negative factors (e.g., predators and disease) (Holling, 1959; Kareiva & Odell, 1987; Huston *et al.*, 1988; Turchin, 1998; Barraquand & Murrell, 2013; Spiegel *et al.*, 2017; Dougherty *et al.*, 2018). Beyond their mechanistic role in driving population/community-level dynamics, encounters are also central to many conservation issues. For instance, an animal’s probability of encountering humans and/or human-related activities is a key indicator of the potential for human-wildlife conflict (Meijaard *et al.*, 2011; Poessel *et al.*, 2013), while encounters with vehicles represent a serious source of mortality for many species (Bennett, 1991; Gibbs & Shriver, 2002; Glista & DeVault, 2008). Additionally, emerging zoonotic diseases pose significant and increasing threats to human health and the global economy (e.g., Rothan & Byrareddy, 2020) and locations of risk are simultaneously viewed as ‘hotspots’ for both conservation and emerging disease (Paige *et al.*, 2015).

Despite the keystone importance of encounter events for a broad range of ecological processes, encounter theory has not kept pace with the developments in animal tracking (Wikelski *et al.*, 2007; Kays *et al.*, 2015; Noonan *et al.*, 2015) or movement modelling (Johnson *et al.*, 2008; Gurarie *et al.*, 2009; Benhamou, 2014; Fleming *et al.*, 2014a; Michelot & Blackwell, 2019; Hooten *et al.*, 2019). Furthermore, existing work has focused primarily on the relationship between animal movement and encounter *rates* (e.g., Gerritsen & Strickler, 1977; Visser & Kiørboe, 2006; Hutchinson & Waser, 2007; Bartumeus *et al.*, 2008; Gurarie & Ovaskainen, 2013; Martinez-Garcia *et al.*, 2020), while no formal framework directly relates individual movement with the spatial *locations* of encounter events in the environment (but see Spiegel *et al.*, 2016). This is a notable limitation as the probability of encountering another individual has well documented effects on animal behaviour. The landscape of fear hypothesis, for example, is based on the concept that prey species alter their behaviour with respect to spatiotemporal predation risk (Lima *et al.*, 1985; Hernández & Laundré, 2005; Kuijper *et al.*, 2013; Laundré *et al.*, 2014). One prediction of this hypothesis is that individuals can devote more time to foraging in areas where the risk of encountering a predator is low, and should therefore balance predation risk against energetic needs while navigating their environments. Without a method for straightforwardly quantifying how the probability of encountering a predator changes in space, however, empirical work has typically relied on intensive field efforts (e.g., van der Merwe & Brown, 2008), and/or *ad hoc* proxy measures to quantify the spatial distribution of predation risk (reviewed in Gallagher *et al.*, 2017).

Beyond the importance of encounters in driving predator-prey dynamics, the way in which intra-specific encounter probability varies in space also shapes socio-spatial arrangements. For example, while many species occupy home ranges with undefended boundaries, others maintain actively defended territories (Powell, 2000). In species with intense and potentially lethal inter-group competition, individuals tend to avoid areas near territorial boundaries, where they are likely to encounter neighbouring individuals (Sillero-Zubiri & Macdonald, 1998; Mech & Harper, 2002; Wrangham *et al.*, 2007a,b), and are more alert when moving through these locations (Wrangham *et al.*, 2007b; Kurihara & Hanya, 2018; Tórrez-Herrera *et al.*, 2020). Indeed, Moorcroft *et al.* (1999) showed how individual coyotes’ (*Canis latrans*) avoidance of areas with a high probability of encountering the scent marks of individuals from neighbouring packs provided a mechanistic basis for the formation of territorial boundaries (see also Moorcroft *et al.*, 2006a). Mechanistic home range approaches, however, require complex, mechanistic models that need to be specifically tailored to each situation, which renders them less generally applicable (e.g., Moorcroft *et al.*, 2006b; Giuggioli *et al.*, 2013). The difficulty in quantifying the location, permeability or even existence of territorial borders (Powell, 2000), has meant that researchers often rely on indirect measures such as patterns of home range overlap when studying animal behaviour at and around territorial boundaries (e.g., Bermejo, 2004; Vander Wal *et al.*, 2014; Tórrez-Herrera *et al.*, 2020).

Although a direct measure of the probability distribution of encounter locations would benefit a wide range of ecological research, there is currently no straightforward method for estimating this from empirical data. Those studies that have attempted to relate individual movement with encounter locations have used randomization approaches to generate null models (Spiegel *et al.*, 2016, 2018), and a general analytical framework is still lacking. Here, we bridge this gap by developing a formal framework for estimating the spatial distribution of encounter events from animal tracking data. To this end, group-living species, by definition, share their home ranges/territories with certain familiar conspecifics while excluding and/or avoiding others (Powell, 2000; Clutton-Brock, 2009). In this context, it may be more informative to consider encounters that occur at the group level. For instance, within-group encounters are likely to occur within the group’s territory/range, and quantifying their locations may provide only marginal insight into the species’ socio-ecology as compared to the locations of encounters *between* groups. We therefore expand our framework beyond the individual-level to allow for the estimation of group-level encounter dynamics. Additionally, most analytical work on encounters model animal movement as either bounded ballistic (Gerritsen & Strickler, 1977; Hutchinson & Waser, 2007), or Brownian (Visser & Kiørboe, 2006; Visser, 2008; Dieker, 2011) motion. A key limitation of these models, however, is that they result in uniformly distributed patterns of space use, meaning encounters between individuals also occur uniformly in space. In stark contrast, most real animals exhibit *non-uniform* space use within spatially restricted home ranges (Burt, 1943; Bowen, 1982; Powell, 2000; Moorcroft *et al.*, 2006b; Kie *et al.*, 2010; Fleming *et al.*, 2014a; Noonan *et al.*, 2019b; Martinez-Garcia *et al.*, 2020), and encounters between individuals do not occur uniformly in space, but are instead concentrated at territorial boundaries (Nievergelt *et al.*, 1998; Bermejo, 2004; Wilson *et al.*, 2012; Ellwood *et al.*, 2017), in/around heavily used habitats and/or habitat-features (Weckel *et al.*, 2006; Whittington *et al.*, 2011), or at key resources (De Boer *et al.*, 2010; Price-Rees *et al.*, 2013). We therefore base our estimation framework on recent analytical work by Martinez-Garcia *et al.* (2020) incorporating non-uniform movement within home ranges into encounter theory.

We first demonstrate the general utility of this method for three simulation-based scenarios that occur routinely in biological systems: i) a population of individuals that occupy relatively distinct home ranges, but with some degree of spatial overlap with neighbours at the edges of their ranges (e.g., Mertl-Millhollen, 1988; Wronski, 2005); ii) a pair of individuals with a hard territorial border between their home ranges (e.g., Henschel & Skinner, 1991; Moorcroft *et al.*, 2006a); and iii) a predator with a large home range that encompassed the home ranges of multiple prey (e.g., Herfindal *et al.*, 2005; Loveridge *et al.*, 2009). We next highlight the real-world applicability of our method on GPS data from white-faced capuchins (*Cebus capucinus*) tracked on Barro Colorado Island, Panama, and sleepy lizards (*Tiliqua rugosa*) tracked in Bundey, South Australia. We show here how the spatial distribution of encounters can provide an estimate of the location of territorial borders, and demonstrate the possibility of gaining insights into species’ ecology by comparing changes in movement patterns against the CDE. We conclude with a discussion of the potential applications of this novel target distribution for better understanding the dynamics of populations of interacting individuals.

## Methods

### Derivation of the individual-level Conditional Distribution of Encounters (CDE)

Before describing the details of our CDE estimator, it is important to note at the outset that although we rely on assumptions in our derivations, the spatial distribution of encounter locations is a target distribution that exists even if the assumptions of the present estimator may not be met by every dataset. Our framework for estimating the spatial distribution of encounter events in the environment is based on the assumption that the mean instantaneous encounter rate 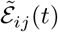 at time *t* between individuals indexed *i* and *j* with location vectors **r**_*i*_(*t*) and **r**_*j*_(*t*) is given by

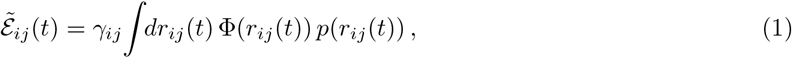

where *γ*_*ij*_ is a proportionality constant that governs encounter risk, also termed the encounter parameter (Gurarie & Ovaskainen, 2013), *r*_*ij*_ = |**r**_*i*_ − **r**_*j*_| is the distance between individuals *i* and *j*, Φ(*r*_*ij*_) is a relative measure of the probability of an encounter occurring when the two individuals are some distance apart (termed the ‘encounter kernel’), and *p*(*r*_*ij*_) is the probability density of separation distance *r*_*ij*_ (Martinez-Garcia *et al.*, 2020). If we assume that encounters are local events, Φ(*r*) limits to a Dirac delta function. For stationary processes (i.e., no change in the means or variances over time), this assumption allows Eq. (1) to be reduced to

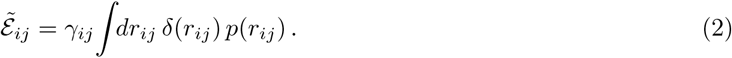

It is important to note that this approximation is valid beyond purely local encounters, provided that the encounter and perceptual ranges are much smaller than the size of the home ranges. Given the individual home range distributions *p*_*i*_(**r**_*i*_) and *p*_*j*_(**r**_*i*_), and the change of variables 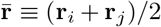 and **r**_*ij*_ = **r**_*i*_ − **r**_*j*_, we then have for a pair of uncorrelated movement processes

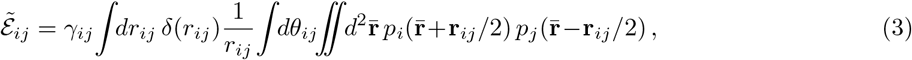

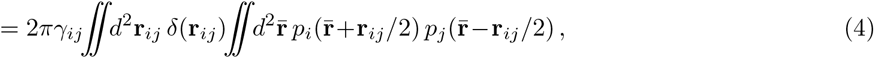

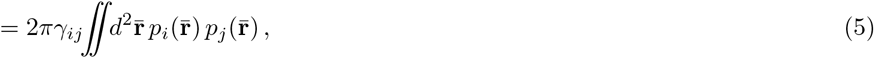

and therefore the contribution to the mean instantaneous encounter rate 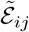 from location **r** is proportional to the product of densities *p*_*i*_(**r**) *p*_*j*_(**r**). Next, because this proportionality is universal across all encounters between individuals *i* and *j*, conditional on an encounter event between *i* and *j* taking place, the probability density function of where that encounter took place must also be proportional to *p*_*i*_(**r**) *p*_*j*_(**r**). Therefore, the CDE is given by the normalized density

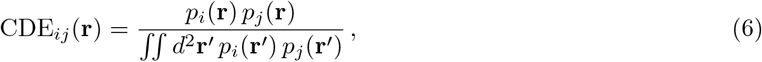

under the assumptions of stationarity, local encounters, and uncorrelated movement. The assumption that movement is uncorrelated across individuals is valid even with cross-correlation-inducing encounters, so long as the individuals’ movement is uncorrelated with one another outside of the encounter event, and the duration of encounters is relatively short compared to the home-range crossing timescales. In Fig. 1 we provide a visualisation of the relationship between movement within home ranges, encounter events, and the CDE for a pair of simulated trajectories generated from uncorrelated Ornstein-Uhlenbeck-Foraging processes (OUF; Fleming *et al.*, 2014a,b), which features autocorrelated positions and velocities, and a defined home range.

**Figure 1:**
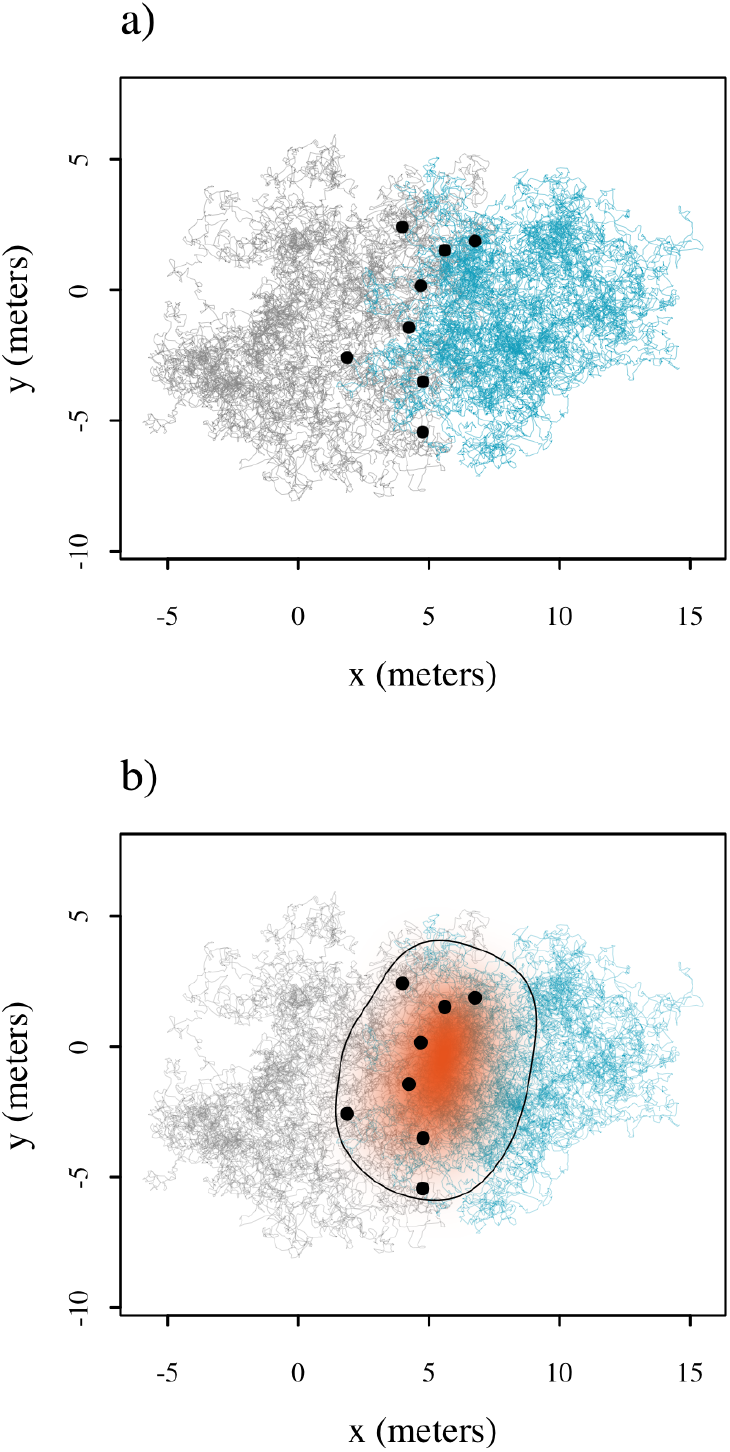
Panel a) depicts the encounter locations (black points) for a pair of trajectories simulated from Ornstein-Uhlenbeck-Foraging (OUF) processes. In b) the resulting CDE estimate (orange shading) and 95% contour are shown.

The lack of a formal framework for estimating the spatial distribution of encounter events has meant that, to date, researchers have often relied on describing patterns of home range overlap as an indirect proxy (e.g., Bermejo, 2004; Vander Wal *et al.*, 2014; Tórrez-Herrera *et al.*, 2020). To place this approach in context with the present work, in Online Appendix S1 we derive an expression for the individual-level CDE in terms of home range overlap via the Bhattachryya Coefficient (BC; Fieberg & Kochanny, 2005; Winner *et al.*, 2018). The Battacharryya Coefficient is a measure of similarity between a pair of distributions that ranges from 0 to 1, 1 if the two distributions are identical, and 0 if there is no shared support (Bhattacharyya, 1943). We find that the denominator of the individual-level CDE in Eq. 6 is given by,

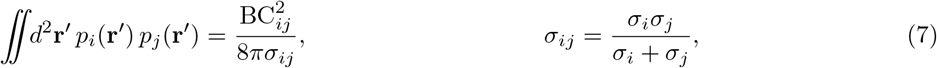

where *σ*_*i*_ and *σ*_*j*_ are the variances of the probability density functions for the position of each individual (both in the *x* and *y* coordinate because movement is assumed isotropic).

### Derivation of the group-level CDE

For group-living species, it may be more informative to consider encounters that occur at the group level (i.e., individuals from one group encountering individuals from another) as opposed to unstructured encounters between all individuals in a population. Here we expand beyond individuals *i* and *j* to consider two groups *I* and *J*, all with otherwise similarly interacting individuals (i.e., no individuals interact more or less than any others), though this latter approximation can be relaxed by a modification of the proportionality constant *γ*_*ij*_. The encounter rate 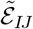 between any of the individuals in group *I* and those in group *J* is given by

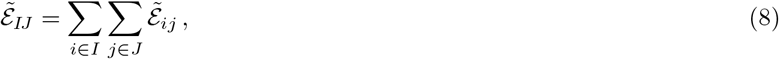

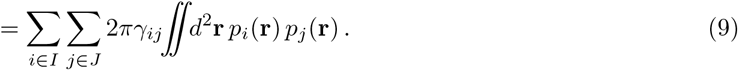

If *γ*_*ij*_ are all similar between groups, then by the same arguments as before, the group-level CDE is given by the normalized density

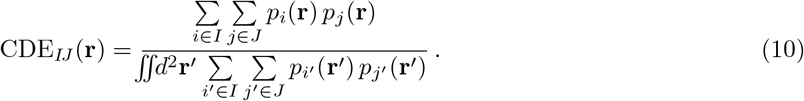

Otherwise, the *γ*_*ij*_, or some quantity proportional to *γ*_*ij*_, must be included in the sums, in which case we have

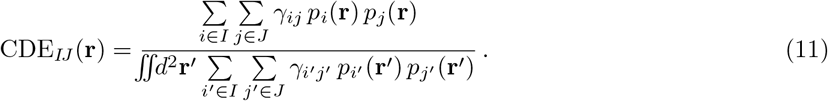

where only the proportional dependence of *γ*_*ij*_ on (*i, j*) matters. That is to say, we care if *γ*_12_ is twice that of *γ*_13_, because it means that individuals 1 and 2 encounter each other twice as much as individuals 1 and 3, all else being equal, but we do not need to know the absolute values of *γ*_*ij*_. We note that the group-level CDE is a function of each individual’s movement, and, as such, makes no overarching assumptions about group dynamics/structure. In other words, the group-level CDE can accommodate groups with varying levels of cohesion, from tightly knit groups that travel as a unit, to more loosely associated fission-fusion groups.

### Confidence intervals for the CDE

Importantly, the CDE is calculated from individual home range estimates, *p*_*i*_(**r**), which are themselves estimates derived from data. CDE estimates should, therefore, be accompanied by a measure of the confidence (Pawitan, 2001). To this end, we propagate uncertainty in the home range estimates into each CDE via a combination of Gaussian reference function (GRF) approximation to the individual home range estimates, followed by the delta approximation (Cox, 2005). Specifically, for each density *p*_*i*_(**r**) we first consider the Gaussian approximation to this density with mean vector ***μ***_*i*_ and covariance matrix ***σ***_*i*_. The individual CDE_*ij*_ is then given by a Gaussian density with mean and covariance

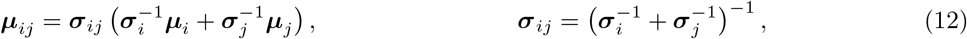

for which we can propagate uncertainty in the mean and covariance parameters via the delta approximation.

I.e., given 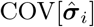 and 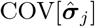, then we have

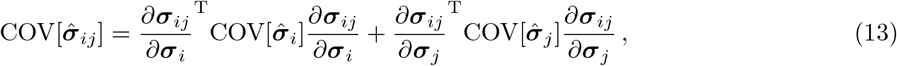

under the assumption of independence in estimates *i* and *j*. With the uncertainty estimates in the Gaussian approximation to the CDE, we then assign that uncertainty to the CDE contours following Fleming *et al.* (2018). The procedure is analogous for the community CDE (Online Appendix S2). This estimator and, accompanying confidence intervals, is fully implemented in the `encounter()` function in the `ctmm R` package (ver. 0.5.11 and later). In Online Appendix S3 we provide details on a simulation study aimed at exploring the statistical performance of our CDE estimator, and the code necessary to reproduce these simulations is provided in Online Appendix S4.

### Workflow for estimating the CDE

Our framework allows researchers to use animal movement data to generate a probabilistic representation of the spatial locations of encounter events in the environment. It does so by incorporating information on individual patterns of home range use into a spatial distribution of encounter events (i.e., the CDE). As modern tracking data are almost invariably autocorrelated (Noonan *et al.*, 2019b, 2020), we built our framework around autocorrelated kernel density estimation (AKDE; Fleming *et al.*, 2015a). We refer readers interested in the statistical efficiency of AKDE to Fleming & Calabrese (2017) and those interested in a comparison of AKDE to other commonly used home range estimators to recent work by Noonan *et al.* (2019b).

With a prepared movement dataset in hand, the first step in the workflow is ensuring that the individuals of interest are range-resident as the CDE is a long-term estimate of encounter location probabilities for movement within home ranges. When the data do not show evidence of range-residency, home range estimation, and therefore CDE estimation, is not appropriate (Calabrese *et al.*, 2016; Fleming & Calabrese, 2017). We therefore strongly recommend starting with visual verification of range-residency via variogram analysis (Fleming *et al.*, 2014a). Once range-residency has been verified, the next step is to fit a series of range-resident continuous-time movement models to the data, such as the Independent and Identically Distributed (IID), Ornstein-Uhlenbeck (OU; Uhlenbeck & Ornstein, 1930), and OU-Foraging (OUF; Fleming *et al.*, 2014a,b) processes. Model selection should then be employed to identify the most appropriate model for the data (Fleming *et al.*, 2014b, 2015b, 2019). With a fitted, selected movement model in hand, AKDE home range estimates can then be estimated (Fleming *et al.*, 2015a; Fleming & Calabrese, 2017; Fleming *et al.*, 2018), and these can be used to obtain CDEs.

While the CDE can be informative on its own, it is also a probability density the has contours just like a home range estimate. Placing contours on the CDE permits the identification of areas where a specified quantile (e.g., 95% or 50%) of encounters will occur, while the error propagation techniques described above enable CIs to be placed on these contours. These CDE estimates may either be the final product of the analysis, or be used in subsequent analyses such as estimating territorial boundaries, identifying changes in behaviour within the 95% CDE area, or habitat features associated with high/low density regions of the CDE. While the workflow we describe involves several steps, the `ctmm` package streamlines this procedure, and a full example of the workflow is shown in Online Appendix S5.

#### Ecologically guided case studies

To demonstrate the ecological significance of the CDE, we generated simulated datasets for three scenarios that routinely occur in biological systems: i) a population of individuals that occupy relatively distinct home ranges, but with some degree of spatial overlap with neighbours at the edges of their ranges (e.g., *Lemur catta* Mertl-Millhollen 1988, *Tragelaphus scriptus* Wronski 2005, *Tiliqua rugosa* Kerr & Bull, 2006a); ii) a pair of individuals with a hard/defended territorial border between their home ranges (e.g., *Crocuta crocuta* Henschel & Skinner 1991, *C. latrans* Moorcroft *et al.*, 2006a); and iii) a predator with a large home range that overlaps the home ranges of multiple prey individuals (e.g., *Lynx lynx Herfindal et al.* 2005, *Panthera leo* Loveridge *et al.*, 2009). Details for these scenarios are presented in turn.

#### Scenario i: Overlapping home ranges with permeable borders

In our first scenario, we simulated tracking data for a population of 7 individuals with equally sized, regularly spaced home ranges. Individuals were arranged hexagonally, For each individual we sampled 1×10^7^ locations from an isotropic, IID process with a spatial variance of 6500 m^2^. We then estimated individual home range areas and the 95% CDE using the workflow described above.

#### Scenario ii: Exclusive home ranges with hard borders

In our second scenario, we simulated tracking data for a pair of individuals with a hard territorial border between mutually exclusive home ranges. For each individual we sampled 5000 locations from IID processes that were Gaussian along the *y*-axis, but half normal along the *x*-axis (i.e., reflected along *x* = 0), and with a spatial variance of 2000 m^2^. As above, we then estimated individual home range areas, and the 95% CDE.

#### Scenario iii: Predator-prey encounters

In our third scenario, we simulated tracking data for a predator with a large home range that encompassed a population of 30 prey individuals. Notably, this scenario would also be appropriate for species where large male home ranges overlap numerous smaller female ranges (Clutton-Brock, 1989; Calabrese *et al.*, 2020). For each individual we sampled 500 locations from an isotropic, IID process. The home range center of the predator was set to (0,0) and the spatial variance to 1×10^7^ m^2^, while the home range centers of the prey were drawn from a bivariate Poisson distribution and spatial variances of 2000 m^2^. We then estimated individual home range areas as above, but here excluded all prey-prey encounters when estimating the CDE, so that this represented only the spatial distribution of predator-prey encounters.

We opted to simulate from IID processes for these case studies as the lack of autocorrelation allowed for more rapid convergence of the estimated home range areas and CDEs (Noonan *et al.*, 2019b). Here again, however, we do not expect any qualitatively different behaviour for data from autocorrelated movement processes. We also note that the amount of data we simulated for each scenario was based on a case-specific trade-off between computation time, accuracy, and visual clarity of the results.

### Empirical studies

#### White-faced capuchins

White-faced capuchins (*C. capucinus*; henceforth capuchins) are New World primates that feed primarily on fruit and invertebrates (Chapman & Fedigan, 1990). They are group-living, with intense (Crofoot, 2007) and potentially lethal (Gros-Louis *et al.*, 2003) inter-group competition. Previous work on this species has found that they have home range areas of ca. 8×10^5^ – 1.5×10^6^ m^2^ (Tórrez-Herrera *et al.*, 2020). Although the perceptual range of *C. capucinus* has not been assessed under field conditions, work on the closely related brown capuchin (*Sapajus apella*) found visual perceptual ranges of ca. 8×10^3^ m^2^ (Janson & Di Bitetti, 1997), satisfying the assumption of perceptual ranges ≪ home ranges. Encounter probability plays an important role in governing capuchin behaviour, and previous work has shown that capuchins tend to avoid areas where they are likely to encounter animals from neighbouring groups (Wrangham *et al.*, 2007a; Tórrez-Herrera *et al.*, 2020), and the location at which neighbouring individuals encounter one another can shape patterns of individual participation in cooperative defense and, ultimately, conflict outcome (Crofoot *et al.*, 2008; Crofoot & Gilby, 2012). We applied our CDE estimation framework to tracking data from five individuals belonging to separate, neighbouring social groups on Barro Colorado Island, Panama.

Locations for these five individuals were collected every 4 minutes during daylight hours (6h-18h) between December 2016 and February 2017, using GPS tracking devices (e-obs GmbH, Gruenwald, Germany). Visual inspection of the tracking data and empirical variograms suggested these individuals occupied fixed (i.e., stationary), effectively distinct home ranges, though with narrow contact zones where home ranges overlapped at inter-group boundaries. Following the workflow described above, we estimated the CDE between these neighbouring groups. We then applied ridge estimation to the estimated CDE using the R package ks (ver. 1.11.7; Duong *et al.*, 2007). Ridge estimation extends the problem of estimating the mode of a unimodal probability density function to higher dimensions (Chen *et al.*, 2015). The resulting ‘density ridges’ are paths that follow the high-density regions of a probability distribution. In the present context, ridges represent locations where encounters are more probable and are therefore likely to be territorial edges. To understand whether the spatial distribution of encounters influenced capuchin behaviour, we subset each individual dataset into movement that occurred inside and outside of the 95% CDE, fit movement models to each of these two subsets, and estimated movement speeds following Noonan *et al.* (2019a). We then compared the movement speeds of animals inside and outside of the 95% CDE using the meta-regression model implemented in the R package metafor (ver. 2.1-0 Viechtbauer, 2010). This approach allowed uncertainty in each individual speed estimate to be propagated into the population level estimate when making comparisons. As a further demonstration of the ecological relevance of the 95% CDE area, we compared the locations of twelve field-observed inter-group encounters, which were not used to estimate the CDE, with the estimated territory boundary and 95% encounter area. To collect these observations, a team of observers conducted on-the-ground observations of the capuchin groups from January 1, 2017 through February 2017. Observations lasted between 5 and 9 hours per day, where observers followed one capuchin group on foot and documented a variety of behaviors. Each of the five capuchin groups were observed at least 3 times during the study period, however two of the groups were observed at a higher frequency (minimum two days per week for each of these two groups). Intergroup encounters between capuchin groups that occurred during the observational time period were documented on an all-occurrence basis and the GPS location of the encounter was recorded using a hand-held Garmin GPS unit.

#### Sleepy lizards

We further applied our workflow to GPS data from three sleepy lizards (*T. rugosa*) tracked in Bundey, South Australia. The sleepy lizard is a large, long-lived, primarily herbivorous, skink from temperate regions of Australia (Kerr & Bull, 2006b). Previous work on this species has found that they have home range areas of ca. 4×104 m2 (Bull & Freake, 1999), and perceptual ranges of ca. 1.3×10^3^ m^2^ (Auburn *et al.*, 2009), again satisfying the assumption of perceptual ranges ≪ home ranges. Visual inspection of the tracking data and empirical variograms suggested these individuals occupied fixed home ranges, satisfying the assumption of stationarity. In contrast to the capuchin example detailed above, sleepy lizards home ranges often overlap extensively with conspecifics (Kerr & Bull, 2006a), but do exhibit some level of territorial defence (Spiegel *et al.*, 2018). GPS locations for these three individuals were collected every 2 minutes over a ~ 2.5 month period in Austral spring 2017 (Oct – Dec). We estimated the CDE for these three animals following the general workflow described above. Because sleepy lizard home ranges overlap extensively with conspecifics, CDE areas with high probabilities may therefore relate more to the location of valuable resources than to territorial boundaries (Leu & Bull, 2016; Sih *et al.*, 2018). To understand how the CDE’s capacity to identify key habitat features compared to more conventional approaches, we contrasted it with the area where the 95% and 50% home range contours of all three individuals intersected.

## Results

### Ecologically guided simulations

In our first scenario of overlapping home ranges with permeable borders (Fig. 2a), we found that this type of socio-spatial arrangement resulted in the 95% CDE area predicting encounters as being more likely to occur in boundary regions and lower probabilities towards home range centers (Fig. 2d). Notably, while the bulk of the probability density was centered on the area of intersection of neighbours’ 95% home ranges, there was support for encounters occurring well beyond these areas and closer to the core of individuals’ home ranges. This highlights how the area of home range intersection can underestimate the area over which encounters are likely to occur, especially when overlap is low (see also Online Appendix S1). In our second scenario for a pair of individuals with a hard territorial border between mutually exclusive home ranges (Fig. 2b), we found that the CDE was correctly distributed along the territorial boundary, but with a spillover that was proportional to the bandwidth of the home range estimates (Fig. 2e). Unsurprisingly, in our predator-prey scenario, 95% of the predator-prey encounters were predicted to occur near the center of the predator’s home range, and the home ranges of prey located near the center of the predator’s home range were entirely within the 95% CDE (Fig. 2c,f). For prey near the periphery of the predator’s home range, however, there were still locations within their home range that were within the 95% CDE, but only in heavily used areas. In other words, predation risk for these individuals was greatest not where they’re closest to the predator’s home range center, but where they spent most of their time relative to where the predator went. This highlights the importance of accounting for non-uniform space use when determining the spatial distribution of predation risk.

**Figure 2:**
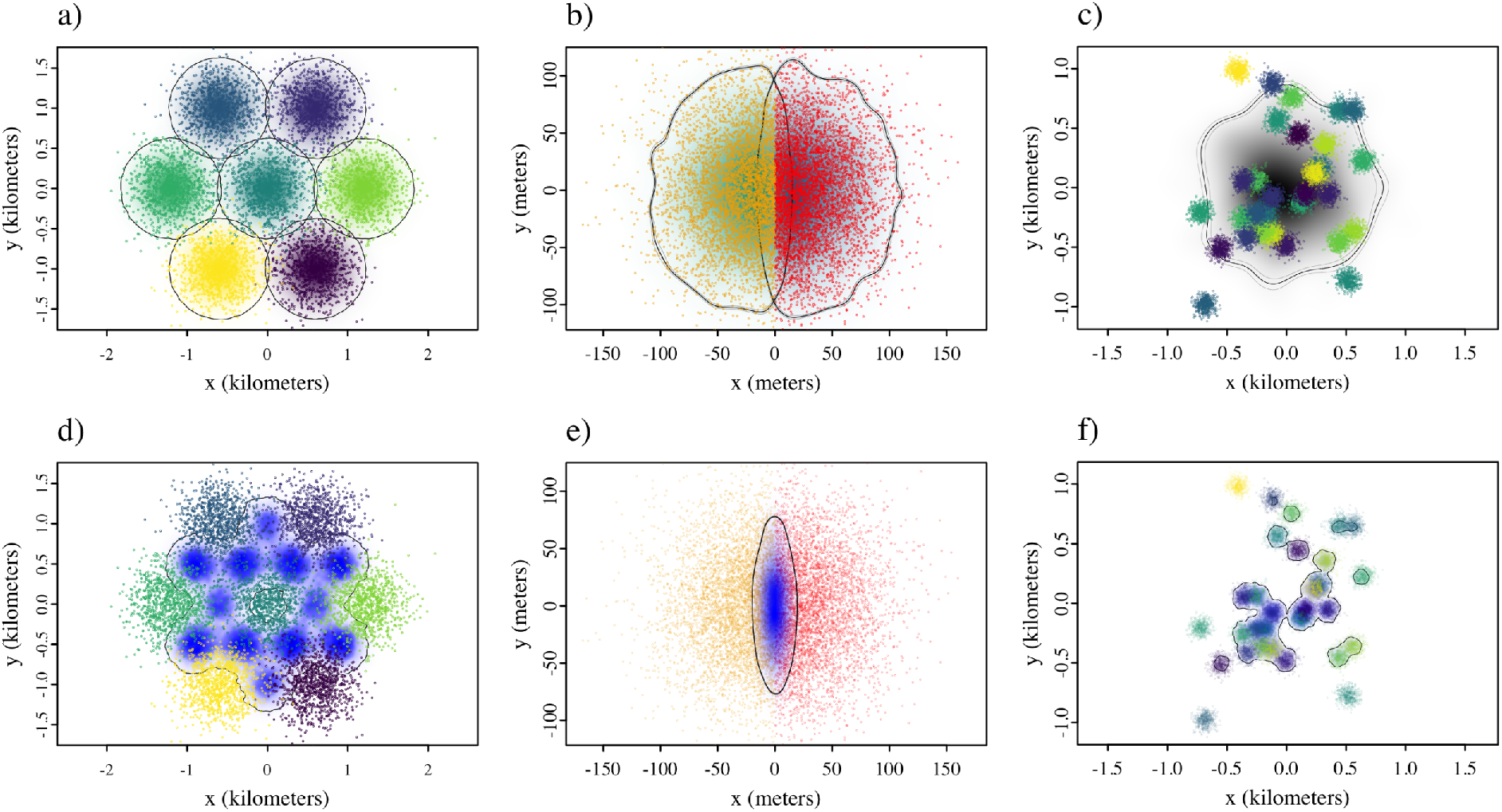
The top row depicts simulated tracking data and home range estimates for: a) a population of individuals that occupy primarily exclusive home ranges but with overlapping boundaries; b) a pair of individuals with a hard territorial boundary between their ranges at *x* = 0; and c) a predator (black density and contours) with a large home range that encompasses multiple smaller prey home ranges (coloured points). In the lower row, panels d), e) and f), the resulting conditional distributions of encounters (CDEs) are shown in blue (darker shadings of blue represent greater probabilities).

### Empirical Case Studies

#### White-faced capuchins

As would be expected for a species with intense inter-group competition, we found that spatial overlap between capuchin home ranges was low (median Bhattacharyya Coefficient: 0.13, range: <0.01–0.28; Fig. 3). Mirroring our simulation based results above (Fig. 2d), the resulting CDE was centered along the boundaries between the individuals’ tracking data, highlighting how most of the inter-group encounters are likely to occur at, or close to, the edges of the individuals’ home ranges. Ridge estimation on the CDE resulted in estimates of the territorial boundaries that mapped onto the edges of each animal’s home range (Fig. 3a). In addition, we found that all 12 directly observed (i.e., by researchers in the field) encounters fell within the 95% contours of the CDE and within 186.6 meters of the estimated territorial boundaries (median 108.7 m; range: 1.7 m — 186.6 m). These findings demonstrate the direct correspondence between the CDE and capuchin ecology.

**Figure 3:**
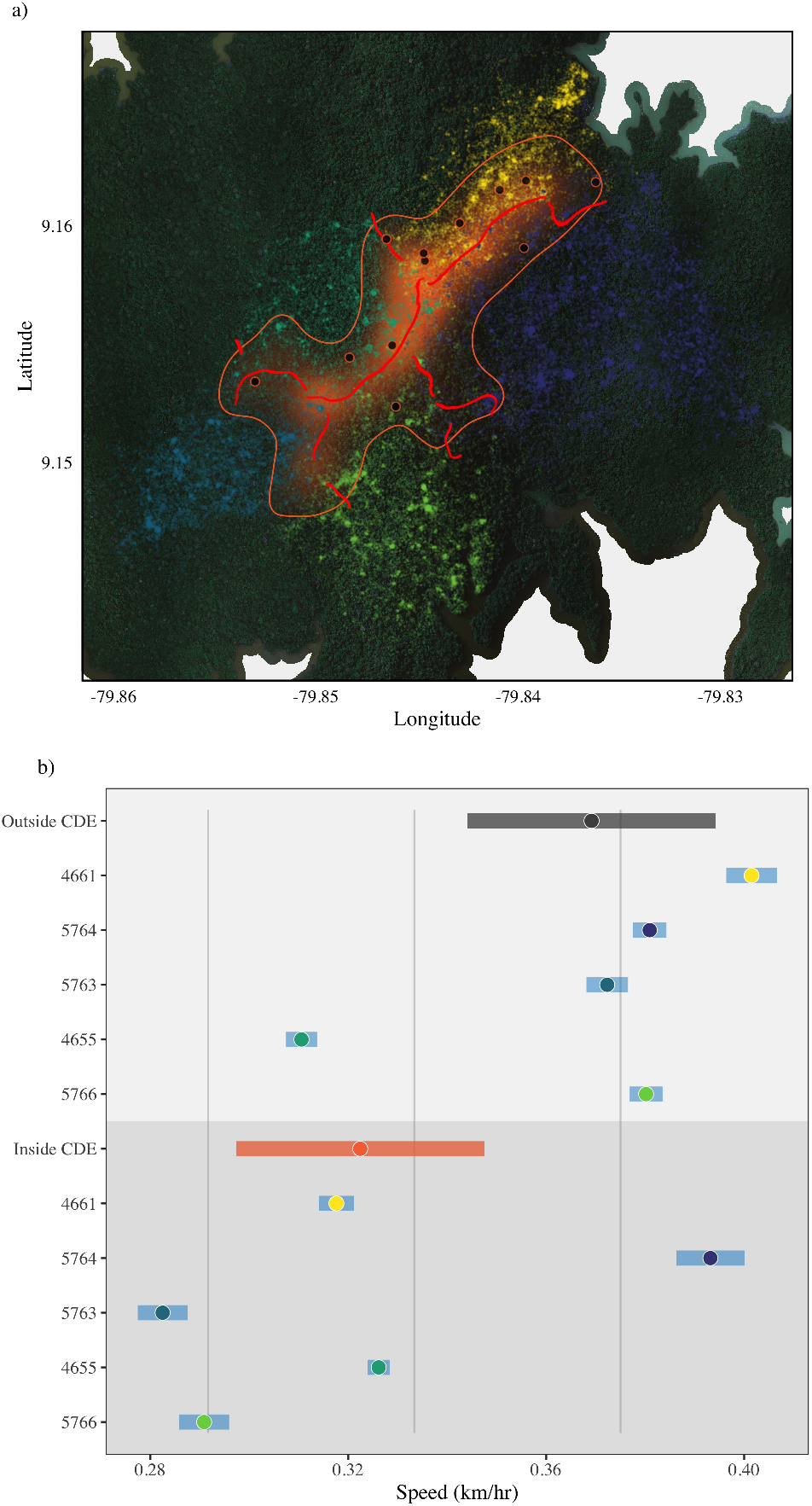
GPS data from five white-faced capuchins (*C. capucinus*) from five neighbouring social groups tracked on Barro Colorado Island, Panama. In panel a) the conditional distribution of encounters (CDE) is depicted in orange shading, while the orange line delineates the 95% contour. The red lines depict the territorial borders estimated via ridge estimation on the CDE, and the large black points represent the locations of field-observed encounters between neighbouring individuals from these five social groups. Note how all of the encounters occurred within the 95% contour of the CDE. In panel b) mean individual and population level speed estimates for movement inside and outside of the 95% CDE are depicted, showing how, on average, animals moved significantly more slowly when in the 95% CDE.

Interestingly, we also found that capuchins modified their movement behaviour in relation to the local probability of encountering a neighbouring individual. While there was no evidence for a general relationship between movement speed and distance from home range centre (R^2^ <0.001, *p* = 0.83), we found that when moving within their home ranges, but not within the 95% CDE, capuchins moved with a mean speed of 0.37 km/h (95% CIs: 0.34 — 0.40). In contrast, animals moved significantly more slowly (*p* = 0.036) when moving through the 95% CDE, with a mean speed of only 0.32 km/h (95% CIs: 0.30 — 0.35; Fig. 3b).

#### Sleepy lizards

We found that although these three animals occupied relatively distinct home ranges with low spatial overlap (median BC: 0.23, range: 0.13 — 0.39), there was a focal point around the only source of standing water in the area where the home ranges of all three individuals intersected (Fig. 4a). For these animals, the bulk of the CDE’s probability density was centered on this water source, suggesting that the majority of encounters are likely to occur at or around this valuable resource (Fig. 4c). If we had applied the conventional approach of describing patterns of home range overlap, however, the area where all three home ranges intersected covered space well beyond the watering hole at the 95% level(Fig. 4b), and there was no location where the 50% contours of all three animals overlap (results not shown).

**Figure 4:**
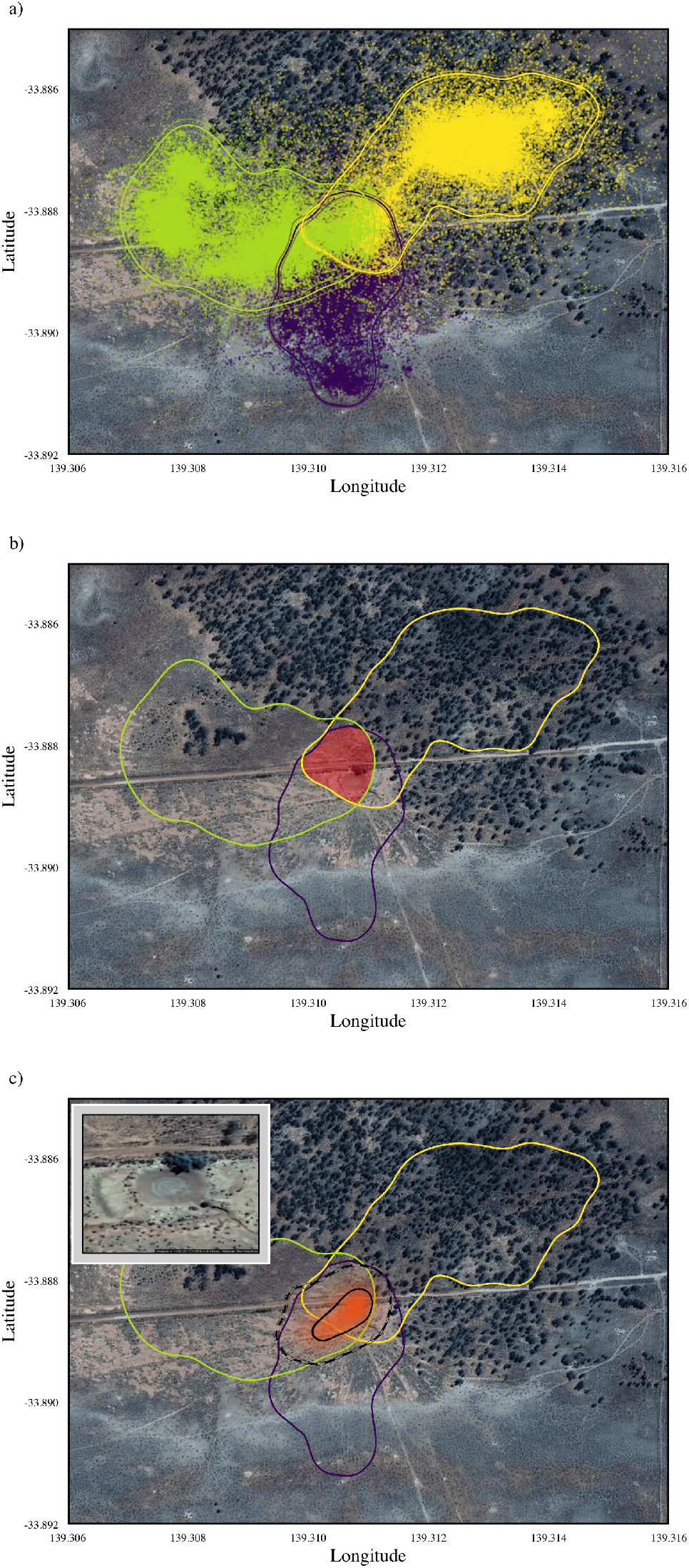
In panel a) GPS data from three sleepy lizards (*T. rugosa*) tracked in Bundey, South Australia are depicted. The contours depict the estimated 95% home range areas ± 95% confidence intervals. Panel b) shows the 95% home range contours, as well as the area where all three 95% home ranges intersect (red shading). In c) the conditional distribution of encounters (CDE) for these animals is shown in orange while the solid black line delineates the 50% contour and the dashed black line delineates the 95% contour. Note how the CDE is centered on the only watering hole in the surrounding area (inset in panel c), while the area of intersection covers space well beyond the watering hole at the 95% level.

## Discussion

Intra- and inter-specific encounters are keystone ecological events that govern the dynamics of many higher-level processes (Holling, 1959; Kareiva & Odell, 1987; Huston *et al.*, 1988; Turchin, 1998; Barraquand & Murrell, 2013; Spiegel *et al.*, 2017; Dougherty *et al.*, 2018). Despite this, the relationship between individual movement strategies and encounter processes has remained conspicuously understudied. Furthermore, previous work has focused almost exclusively on relating animal movement to encounter *rates* (e.g., Gerritsen & Strickler, 1977; Visser & Kiørboe, 2006; Hutchinson & Waser, 2007; Bartumeus *et al.*, 2008; Gurarie & Ovaskainen, 2013; Martinez-Garcia *et al.*, 2020). This has left researchers interested in understanding the spatial dynamics of encounter processes with only empirically based null models (Spiegel *et al.*, 2016, 2018) or indirect *ad hoc* measures of the spatial distribution of encounters. In response, we have derived an estimator of the spatial distribution of encounter events that builds straightforwardly off of one of the most ubiquitous analyses in movement ecology – home range estimation.

### Properties and assumptions of the CDE

Before discussing the properties of the CDE estimator described in the present work, it is crucial to note that the spatial distribution of encounter locations exists even if the assumptions of our estimator are not met by the data. In other words, although the present estimator may not be appropriate for every dataset, it does not negate the existence of the CDE. In deriving our CDE estimator, we relied on three key assumptions to maintain analytical tractability: i) stationarity in the movement processes; ii) that encounters are local events; iii) that movement is uncorrelated across individuals. Stationarity in this context refers to the fact that we are assuming the individuals of interest are range-resident, and do not exhibit a range shift, or major change in home-range behaviour over time. While large-scale analyses suggest that this assumption holds true for many animal tracking datasets (Noonan *et al.*, 2019b, 2020), we recommend verification prior to analysis, as significant changes in movement behaviour (e.g., range shifts, migrations, dispersals, etc.) will clearly influence the area over which encounters occur. In terms of encounters being local events, we anticipate this assumption holding for many species as encounters tend to occur over much shorter distances than the radii of their home range areas (e.g., Muirhead & Sprules, 2003; Middleton *et al.*, 2013). For species with large perceptual ranges, however, the encounter kernel in Eq. (1) can be carried through in the derivations and modified to account for the greater area over which an encounter can be considered to happen. Perhaps the most important assumption of the present framework is that movement is uncorrelated across individuals. As noted above, the assumption of uncorrelated movement is still valid with cross-correlation-inducing encounters, so long as the individuals’ movement is uncorrelated outside of the encounter event, and the duration of encounters is relatively short compared to the home-range crossing timescales (e.g., encounters on the order of minutes vs. range crossing times of days). Nonetheless, correlated movement is a well documented phenomenon that is likely to occur in a wide range of species (Couzin *et al.*, 2005; Strandburg-Peshkin *et al.*, 2015; Calabrese *et al.*, 2018). While expanding the current framework to account for cross-correlated movement was beyond the scope of the present study, future work on this topic is clearly warranted.

In terms of accuracy, our simulation study revealed how, because the CDE is estimated conditionally on multiple home range estimates, any biases in these will be propagated into CDE estimates. Accurate home range estimates are therefore critical for the CDE to accurately reflect the spatial distribution of encounters between tracked individuals (see also Winner *et al.*, 2018). For a discussion on how to obtain accurate home range estimates, we refer readers to (Noonan *et al.*, 2019b). In addition, more effort remains to derive bias corrections for both the Gaussian and kernel estimates. As a further limitation, it is crucial to note that the CDE requires multiple individuals, of potentially different species, to be tracked in the same place at the same time, and provides no information on encounters with unmonitored animals. In other words, if the CDE has a low location-specific probability, this does not necessarily mean an encounter is unlikely if an individual is moving through an area that is regularly visited by untracked animals. Good coverage of the local population is therefore necessary for the CDE to fully capture the spatial distribution of encounters. Importantly, while data density can be a limiting factor in practical applications, this fundamental limitation also exists for any method that quantifies encounter processes, including randomizing paths to generate null models (Spiegel *et al.*, 2016, 2018), comparing home range overlap (e.g., Bermejo, 2004; Vander Wal *et al.*, 2014; Tórrez-Herrera *et al.*, 2020), or applying mechanistic home range analysis (Moorcroft *et al.*, 1999). We therefore recommend that researchers interested in understanding encounter dynamics focus their data collection on good coverage of a localised population.

### The ecological importance of encounter distributions

Our empirical case studies show how the CDE can be used to straightforwardly quantify hitherto intractable aspects of population/community dynamics. For instance, while many truly territorial species actively defend borders (Stamps & Buechner, 1985; Powell, 2000) for other species, borders tend not to be rigid territorial boundaries, but permeable contact zones (e.g., Stewart *et al.*, 1997; Anich *et al.*, 2009; Ellwood *et al.*, 2017). Understanding behaviour at and around territorial boundaries, however, is a deceptively challenging question that, to date, has relied on labor intensive field efforts (e.g., Kruuk, 1972; Delahay *et al.*, 2000; Kilshaw *et al.*, 2009) or modelling species-specific mechanistic processes (Moorcroft *et al.*, 1999; Giuggioli *et al.*, 2013). In the capuchin data we analysed, we showed how the 95% CDE provided an accurate predictor of twelve inter-group encounters that were field-observed independent of the tracking data, as well as how application of ridge estimation on the CDE yielded an objective estimate of the territorial boundaries. Additionally, the CDE can be used to identify any changes in movement behaviour that directly result from location-specific encounter probability. For instance, despite a lack of evidence for a general relationship between movement speed and distance from home range centre, we found that when capuchins moved through the area contained within the 95% CDE, they did so at a significantly slower speed than when moving through areas where encounters with neighbours were low. While the mechanisms behind this difference were not explored in the present study, this agrees with previous work which found that capuchins exploit resources in inter-group boundary areas more thoroughly, and spend longer feeding in each patch they encounter (Tórrez-Herrera *et al.*, 2020).

Application of our CDE framework can also be used to identify key resources, and quantify location-specific potential for competition. In species with high spatial overlap, areas where encounters are more likely to occur probably relate more to valuable resources than territorial boundary dynamics. For example, in our analysis of the sleepy lizard data, instead of reflecting patterns of territoriality, the CDE was localised around the only source of standing water in the three animals’ home ranges. This finding agrees with previous work in this study system that found that this water source was a valuable resource that influences the population’s spatial ecology (Leu & Bull, 2016). Notably, in this respect, neither the area of intersection of the 95% nor the 50% home ranges for these animals adequately identified this aspect of these sleepy lizard’s space use. This highlights how the CDE directly captures ecologically relevant information. In contrast, describing patterns of home range overlap can require researchers to make some level of subjective judgment in their analyses, due in large part to the fact that this approach fails to account for non-uniform patterns in space use and encounter probability. Beyond the utility of the CDE in our empirical case studies, this framework can be used to investigate a wide range of intra- and inter-specific relationships such as predator-prey and/or community dynamics, understanding how encounter rates vary in corridors, along key migration routes, in larger vs. smaller reserves, etc. At the intra-specific level, this method can quantify variation in individual sociability and hot-spots of encounters, augmenting many social network studies that focus on individual heterogeneity and neglect the spatial component of social interactions (see also Spiegel *et al.*, 2016).

### The CDE as a conservation tool

Although the case studies presented here were focused on demonstrating the utility of the CDE for understanding basic ecological processes, understanding where key types of encounters happen is also valuable from a conservation perspective. Human-wildlife conflict represents a major conservation concern which, over the past 20 years, has gone from a barely recognized issue to a major conservation focus (Distefano, 2005; Dickman, 2010). The probability of encountering humans and human-related activities is a key indicator of the potential for human-wildlife conflict. Here, the CDE can be extended to model and estimate the distribution of conflict events and help focus management actions and interventions where they will have maximum effect. Road traffic incidents, for instance, where an animal encounters and is potentially hit by a car, represent a serious source of mortality for many species (Bennett, 1991; Gibbs & Shriver, 2002; Glista & DeVault, 2008), and carry an economic cost of ~$1 billon/year in Europe (Bruinderink & Hazebroek, 1996) and ~$8 billion/year in the United states (Huijser *et al.*, 2017). Given that researchers routinely use tracking data to study interactions between animals and vehicles (Neumann *et al.*, 2012; Zimmermann *et al.*, 2014; Murray & St. Clair, 2015), application of the CDE in this context would aid in better understanding how variation in traffic volume and animal movement affects road traffic incidents risk, and would provide crucial baseline information for developing effective mitigation strategies. Additionally, emerging zoonotic and anthroponotic diseases pose significant and increasing threats to both human and animal health respectively (Estrada *et al.*, 2017; Rothan & Byrareddy, 2020), and locations of risk are simultaneously viewed as ‘hotspots’ for both conservation and emerging disease (Paige *et al.*, 2015). Given the importance of accounting for animal movement when modeling disease dynamics (Dougherty *et al.*, 2018), the CDE represents a potentially fruitful tool for identifying disease transmission hotspots.

### Concluding remarks

In this study we have introduced a new theoretical concept, the conditional distribution of encounter events. Conceptually, the CDE describes how encounters change in space for movement within home ranges, and solidifies what has heretofore only abstractly been defined. Furthermore, we have derived this distribution and confidence intervals, implemented its statistical estimator for empirical movement data, and demonstrated the broad ecological relevance of the CDE. Notably, the general estimation framework developed in this work builds straightforwardly off of home range estimation, and, as such, requires no specialised data collection protocols. CDE estimation thus allows researchers to quantify hitherto intractable aspects of population/community dynamics without the need for intensive field efforts, complex data collection, or relying on *ad hoc* indeces. This method is now openly available via command line interface through the `ctmm R` package (Calabrese *et al.*, 2016; CTMM Initiative *et al.*, 2019) or through the web-based graphical user interface available at ctmm.shinyapps.io/ctmmweb/ (Dong *et al.*, 2017; Calabrese *et al.*, 2020).

## Appendix S1 Relationship between the CDE and home-range overlap

The lack of a formal framework for estimating location specific encounter probabilities has meant that, to date, researchers have had to rely on describing patterns of home range overlap as a proxy for the spatial distribution of encounter events (e.g., Bermejo, 2004; Vander Wal *et al.*, 2014; Tórrez-Herrera *et al.*, 2020). To place this approach in context with the present work, in the appendix we obtain an expression for the individual-level CDE in terms of pairwise home range overlap via the Bhattachryya Coefficient (BC; Fieberg & Kochanny, 2005; Winner *et al.*, 2018).

We first rewrite the definition of the individual-level CDE of Eq. (6) using the definition of the inner product of the PDF for the position of the individuals *p*_*i*_ · *p*_*j*_ = *∫ d****r′*** *p*_*i*_(***r′***)*p*_*j*_(***r′***),

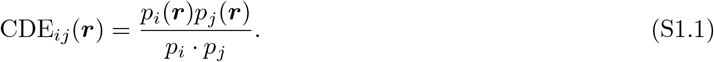

If we assume that individual movement is isotropic and described by a pair of Ornstein-Uhlenbeck (OU) processes, then we can use the results obtained in recent work by Martinez-Garcia *et al.* (2020). Specifically, Eq. (16) in Martinez-Garcia *et al.* (2020) relates the mean instantaneous encounter rate, 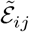, to the BC,

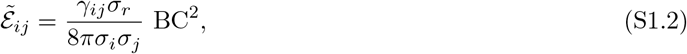

and Eq. (17) in the same work links the mean instantaneous encounter rate with the inner product of *p*_*i*_ and *p*_*j*_,

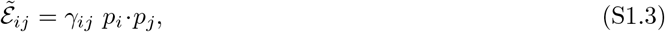

where *σ*_*i*_ and *σ*_*j*_ are the variances of the probability density functions (PDFs) for the position of each individual (both in the *x* and *y* coordinate because movement is assumed isotropic), and *σ*_*r*_ ≡ *σ*_*i*_ + *σ*_*j*_ is the variance of the difference between individual positions. Equating Eqs. (S1.2) and (S1.3), we obtain the inner product as a function of the BC,

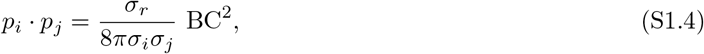

which can be inserted into Eq. S1.1

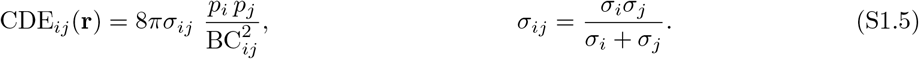

Inserting the leftmost Eq. S1.5 into Eq. 6, we find that the denominator of the individual-level CDE in Eq. 6 is given by

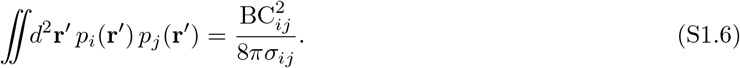

## Appendix S2 Community Gaussian reference function approximations

In this appendix we provide details on propagating uncertainty in the community level home-range estimates to the community-level CDE.

## 2.1 Community GRF approximation

For the community-level Gaussian reference function (GRF) approximation, first we simplify the GRF CDE to a weighted sum of normalized Gaussian functions

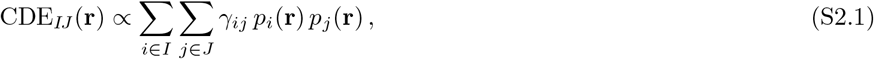

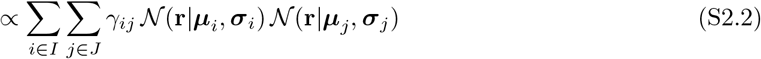

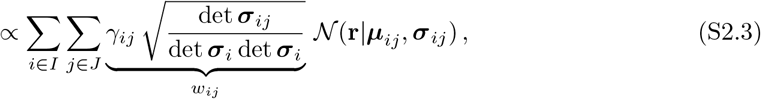

with weights *w*_*ij*_. The first and second moments of the CDE are then given by the corresponding weighted averages, so that the first two cumulants are

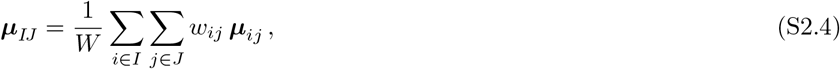

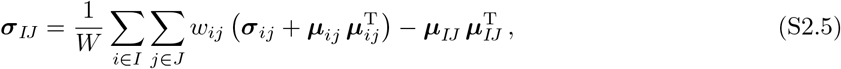

where *W* = Σ_*i*∈*I*_ Σ_*j*∈*J*_ *w*_*ij*_.

## Appendix S3 Statistical Performance of the CDE Estimator

In this appendix we derive a general expression for the individual-level conditional distribution of encounters (CDE) for a pair of Ornstein-Uhlenbeck processes. We selected the OU process here as it benefits from being mathematically tractable, while also featuring non-uniform space use and applying frequently in practice (Noonan *et al.*, 2019b). This also provided a link between this study and Martinez-Garcia *et al.* (2020) that was also based on OU processes, and thus between encounter locations and encounter rates. This derivation allowed us to establish exact expressions that estimated CDEs could be compared against. Using this expression, we performed a simulation study aimed at exploring the statistical efficiency and asymptotic consistency of our CDE estimation framework.

### The Ornstein-Uhlenbeck model

We start by considering a pair of two-dimensional, independent (i.e., without cross-correlations between the individual trajectories) Ornstein-Uhlenbeck processes. The position of each individual evolves in time according to a pair of independent stochastic differential equations,

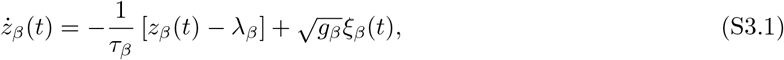

where the subscript *β* indicates each of the two coordinates in the 2D space: *β* ∈ {*x, y*}, *z* is the location of the individual and the dot indicates a time derivative, (*λ*_*x*_, *λ*_*y*_) gives the home range center, and (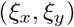) is a mean-zero and unit-variance Gaussian white noise process with units of time^−1/2^. Importantly, because we are going to consider the limit of purely-local perception, *q* = 0, we can consider non-isotropic movement and thus both ***τ*** and ***g*** are 2D vectors. The parameter ***τ*** provides a metric for home-range affinity and ***g*** modulates the size of the home range via the intensity of the stochastic contribution to the movement of the individual. Because the OU model is a Gaussian process, the *i*-th individual-position probability density function (PDF), *p*_*i*_(***r***; *t*), is completely determined by its mean value, ***μ*** = (*μ*_*x*_, *μ*_*y*_), and covariance matrix, Σ. For simplicity, we will consider OU models that are in the stationary regime (the initial condition is drawn from a Normal distribution with moments equal to the stationary moments of the OU model) and diagonal covariance matrices. Under these assumptions the PDF for the position of individual *i* is time-independent, *p*_*i*_(***r***), and completely defined by the mean and the variance of each component of the movement,

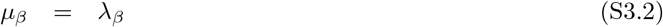

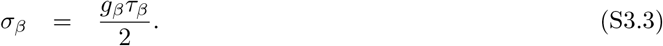

### General expression for the individual-level CDE

We are interested in obtaining the individual-level Conditional Distribution of Encounters (CDE), that is, the PDF for the location of the encounters between two individuals, *i* and *j*, that move according to the OU models described above. Following Martinez-Garcia *et al.* (2020), we assume that the home-range centers of both individuals are at a distance 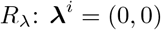 and 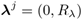^1^. The CDE is proportional to *p*_*i*_(***r***)*p*_*j*_(***r***)

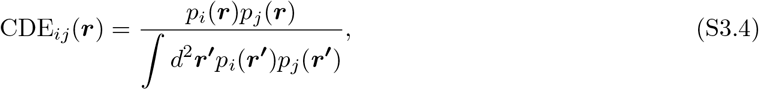

notice that the normalization factor is proportional to the encounter rate in the local-perception limit as derived in Martinez-Garcia *et al.* (2020). Solving the integral for the normalization in Eq. (S3.4), we obtain the CDE,

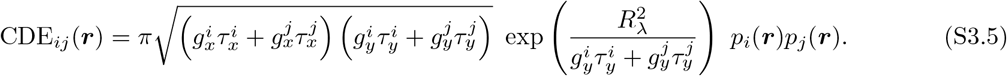

In Figure S3.1, we provide a visual comparison the individual-level CDE as obtained in Eq. (S3.5) with the one reconstructed from encounters between two simulated OU trajectories. If individual movement is isotropic, the expression for the individual-level CDE reduces to

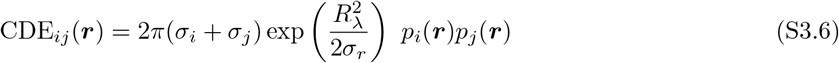

**Figure S3.1:**
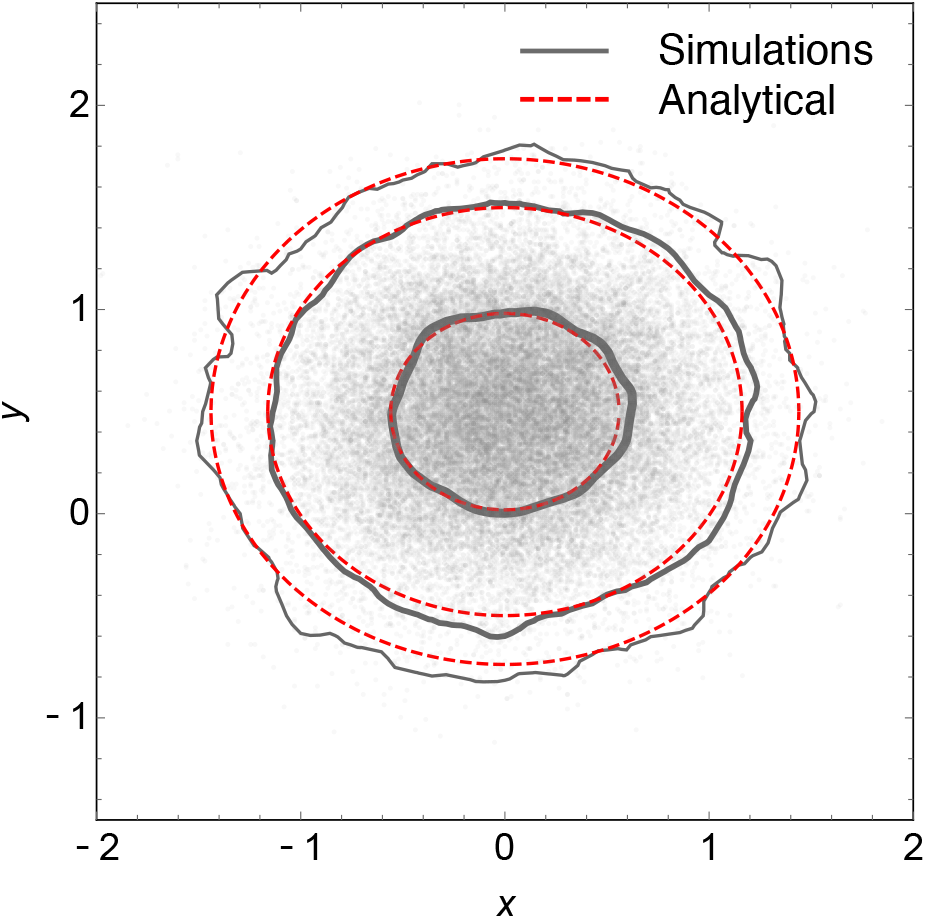
The contours indicate 50%, 95% and 99% of the probability mass. Gray contours correspond to simulations and red dashed, to the analytical result Eq. (S3.5). The gray dots indicate the location of all the 42142 encounters recorded during the 10^7^ time units simulation. Parameter values: *R*_λ_ = 3, 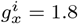, 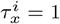, 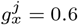, 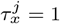, 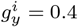, 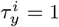, 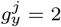, 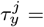, 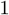 *t* = 10^7^.

### Estimator performance study

Using the general expression for the individual-level CDE derived above, we performed a simulation study aimed at exploring the statistical efficiency and asymptotic convergence of our CDE estimation framework.

Tracking data were simulated from a pair of isotropic OU processes. For these simulations, we set the home range crossing timescales (*τ*_*i*_ and *τ*_*j*_) to 1 day, the spatial variances (*σ*_*i*_ and *σ*_*j*_) to 51,840m^2^, and the home range centers to *μ*_*i*_ = (0m,0m), and *μ*_*j*_ = (0m,800m). We note that although we explored only a single parameter regime, previous work suggests that other parameterisations of these models should result in qualitatively similar behaviour (Winner *et al.*, 2018; Noonan *et al.*, 2019b; Fleming *et al.*, 2019). To confirm asymptotic convergence, we sampled from these processes at a frequency of 8 locations/day and manipulated the sampling duration from 4 to 4096 days in a doubling series. We examined only a single sampling frequency as AKDE can accommodate dataset-specific autocorrelation structures, and such manipulations would not have impacted the underlying home range estimates (see Winner *et al.*, 2018; Noonan *et al.*, 2019b). For each of these pairs of simulated datasets we estimated the CDE following the workflow described above. We then compared the 95% area of the estimated CDE with the 95% area of the true CDE, to quantify how closely the two agreed. The simulation and estimation process was then repeated 1000 times for each sampling duration and statistical performance was evaluated by examining the mean accuracy over replicates. Simulations were performed in the R environment, using the ctmm package and the code necessary to reproduce these simulations is provided in Online Appendix S5.

### Simulation results

Under the parameter and sampling regimes we tested, we found that our CDE estimator tended to exhibit bias for very short sampling durations, but with asymptotic convergence in the large sample size limit (Fig. S3.2).

This bias at short sampling durations was driven by small-sample-size bias in each of the individual home range estimates propagating into the CDE estimate. Importantly, however, we also observed an appropriate bias-variance tradeoff at small sample sizes (i.e., as accuracy decreased the variance increased). We also note that, all else being equal, convergence should happen more rapidly as the home range crossing time becomes shorter, and vice versa, as home range crossing time directly underlies the effective sample size for home range estimation.

As can be seen in the Gaussian results (red line in Fig. S3.2), the remaining bias is due to negative bias in the underlying home range estimates, which is common for downstream analyses that are conditional on multiple home-range estimates (Winner *et al.*, 2018). Because the remaining biases result primarily from the individual range estimates, optimizing the individual bandwidths for the CDE, and deriving a bias correction akin to the area correction derived by (Fleming & Calabrese, 2017) may provide a means of further reducing bias in the CDE, but this was beyond the scope of the present study.

**Figure S3.2:**
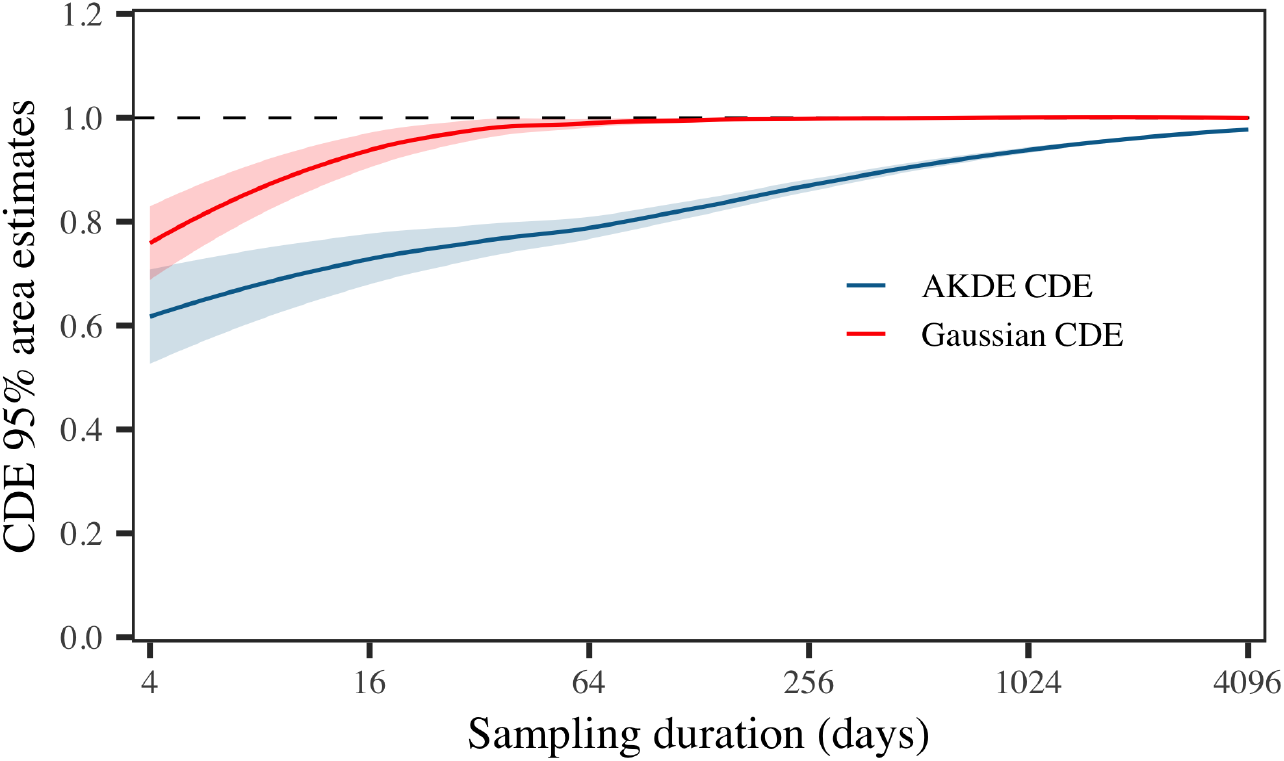
Figure depicting the results of simulations exploring the accuracy of the CDE estimator based on Gaussian home ranges (red) and AKDE home ranges (blue). The solid lines depicts the mean ratio between the estimated and the analytical CDEs, and the shaded areas depict the variance around the mean. The dashed line at *y* = 1 denotes the true value the estimates should be converging to.

## Appendix S4 R script for reproducing the simulations

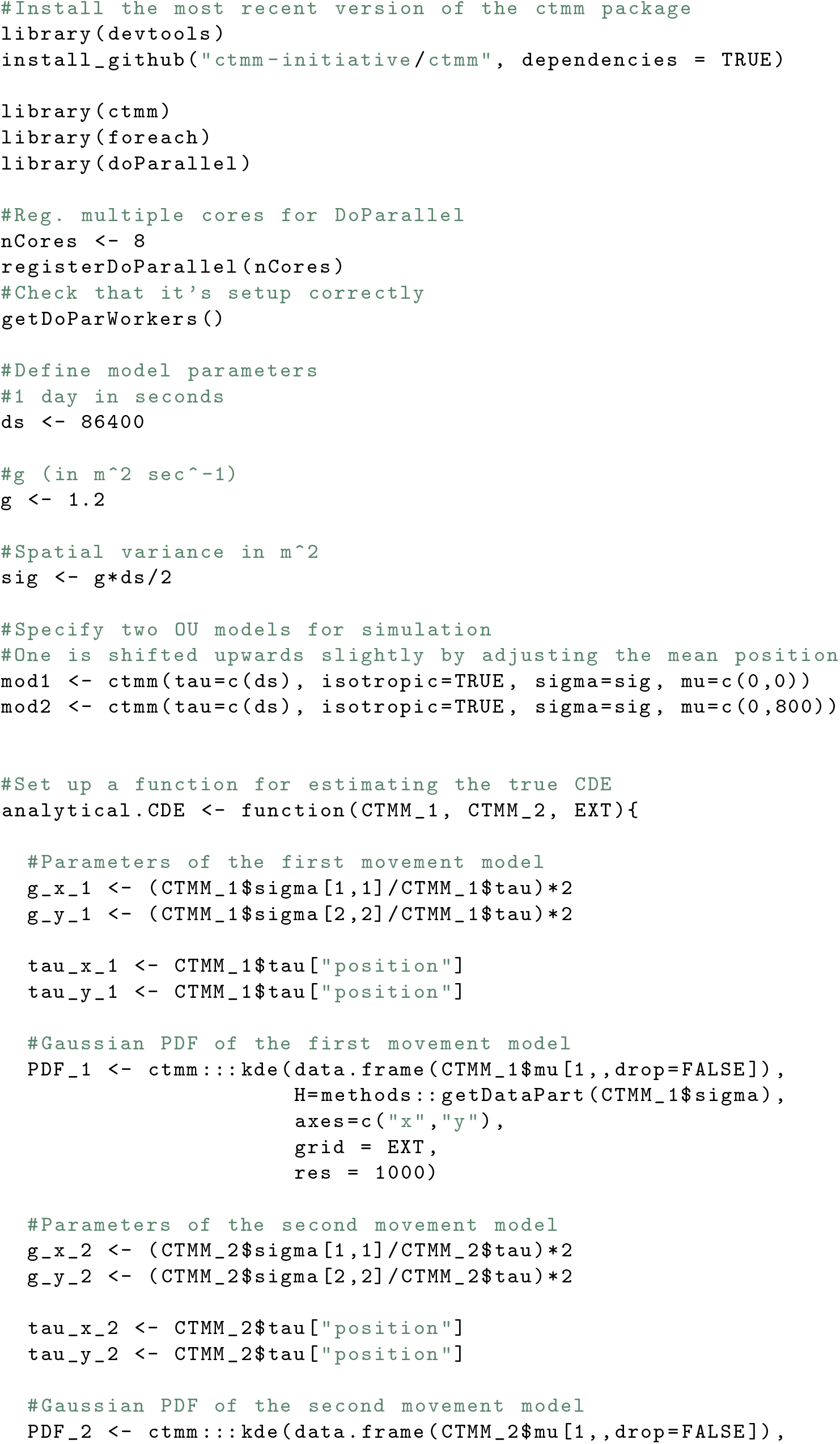
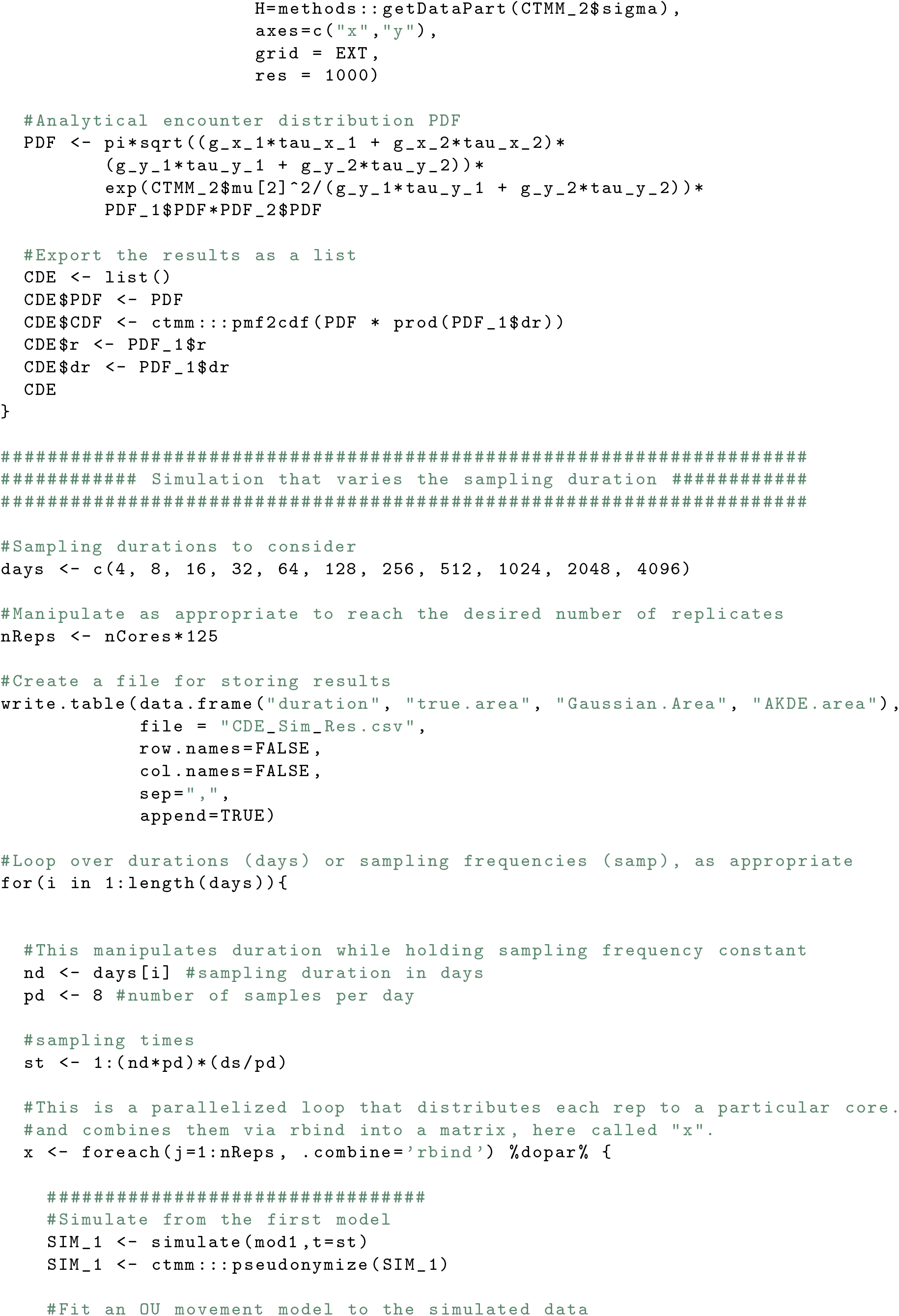
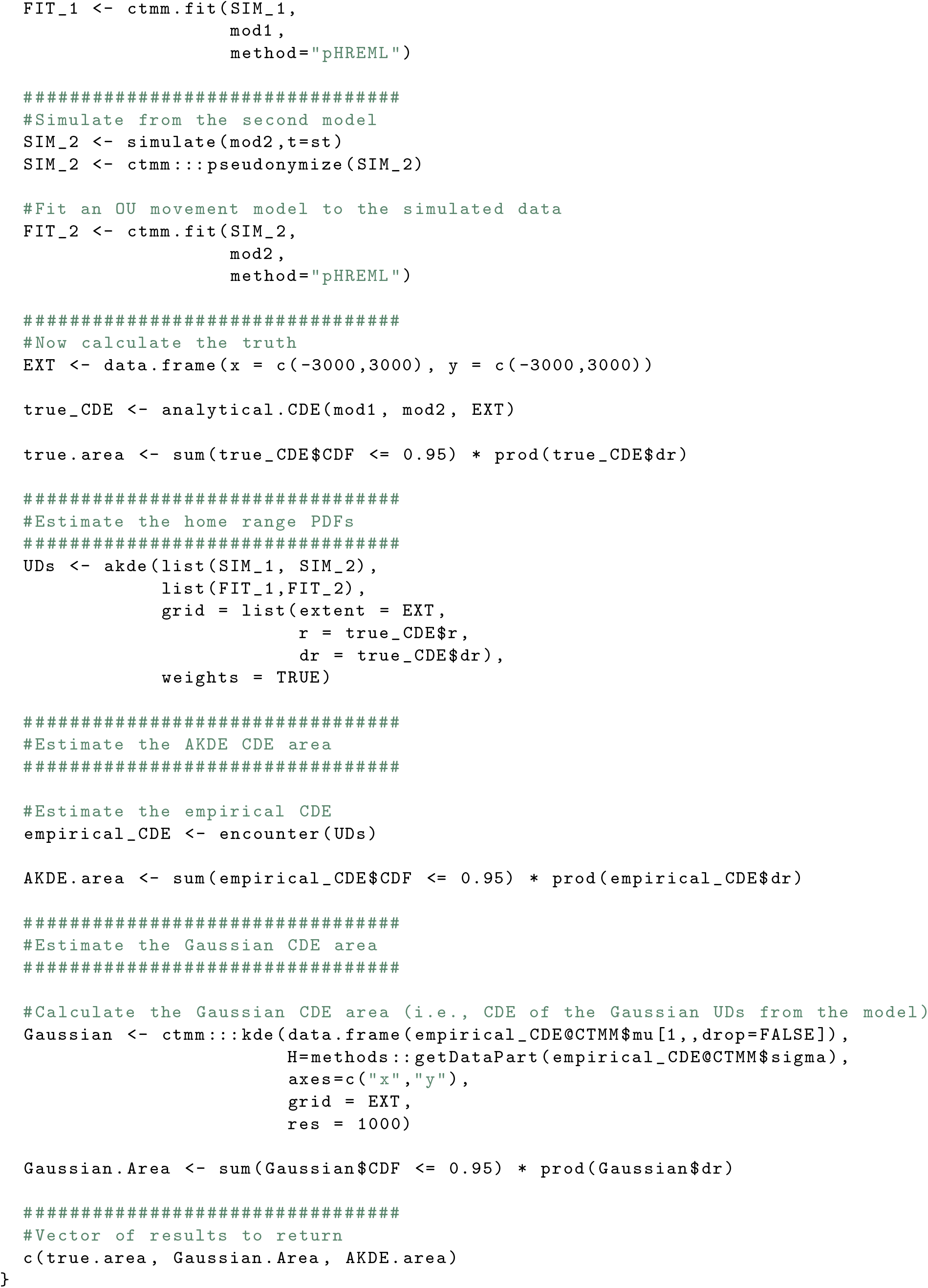
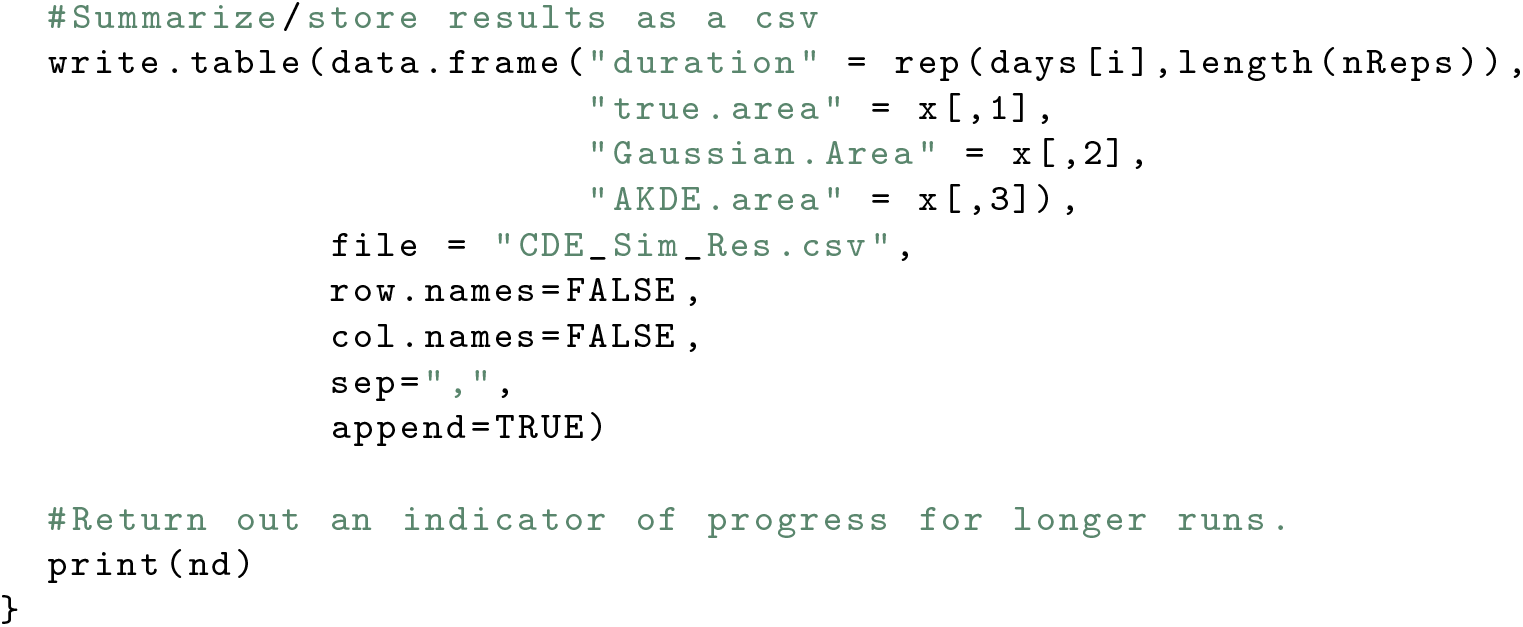

## Appendix S5 Workflow for estimating the Conditional Distribution of Encounters in ctmm

In this appendix we provide a worked example of how to estimate the Conditional Distribution of Encounters (CDE) detailed in the main text. This is carried out on GPS data from two African buffalo (*syncerus caffer*) tracked in Kruger National Park, South Africa (Getz *et al.*, 2007; Cross *et al.*, 2016) These data are openly available within the `ctmm` package.

### Step 1: Data import and confirmation of range-residency

As detailed in the main text, the first step of CDE estimation after data import is to ensure that the individuals of interest are range-resident. We therefore strongly recommend starting with visual verification of range-residency via variogram analysis (Fleming *et al.*, 2014a). In `ctmm` this is carried out by visually inspecting the output of the variogram function.

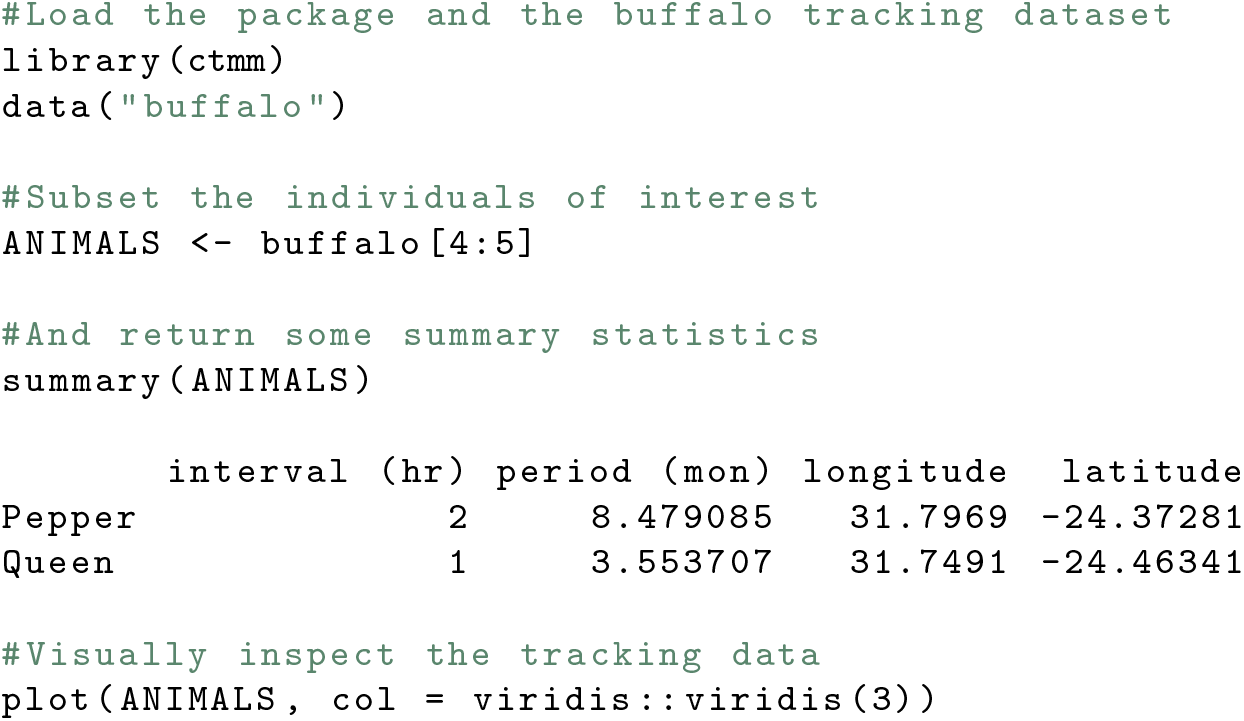

**Figure S5.1:**
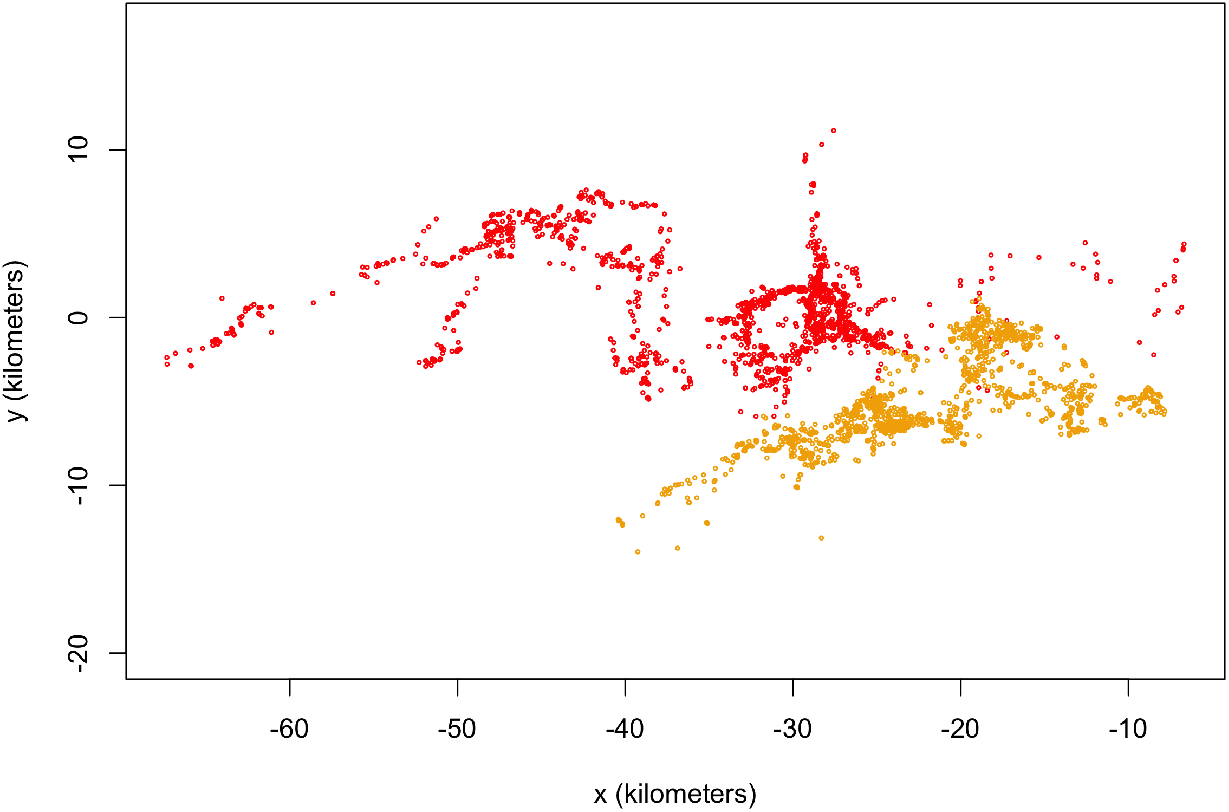
A scatterplot of the GPS positional observations for two African buffalo (*syncerus caffer*) tracked in Kruger National Park, South Africa.

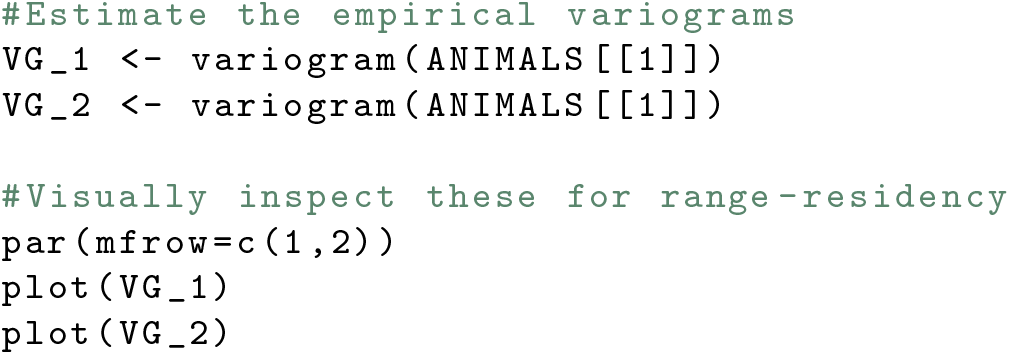

**Figure S5.2:**
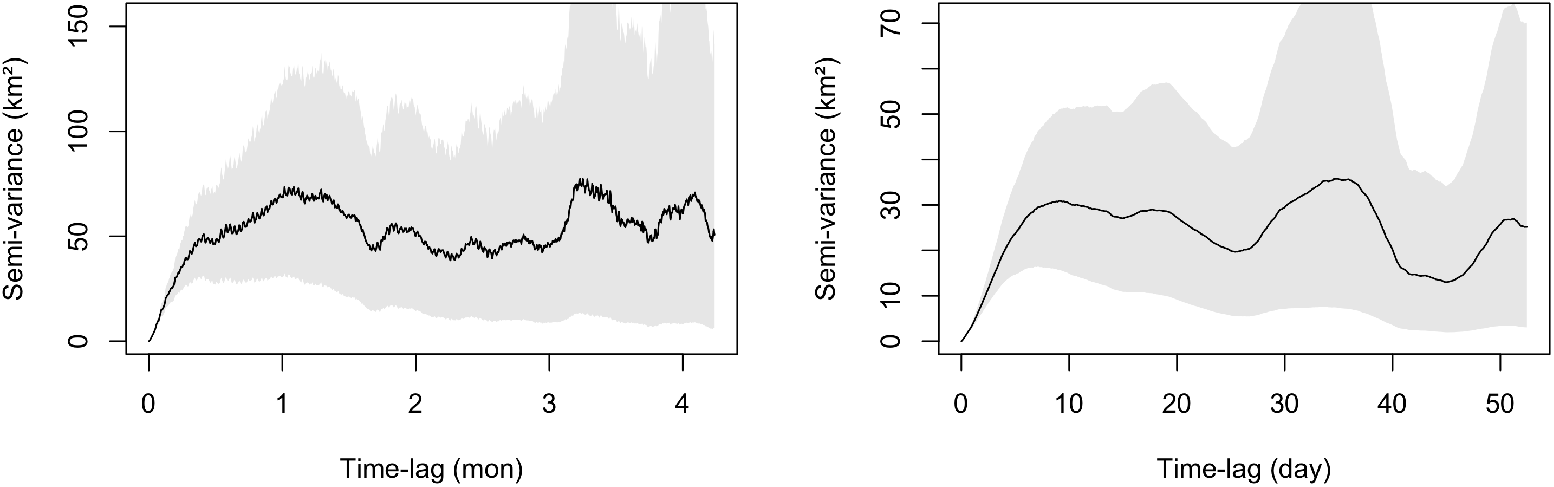
Empirical variograms estimated from the two individuals’ tracking data. Note how both variograms have a clear asymptote, which demonstrates space use is not infinitely diffusive over time, but rather restricted to a home range.

### Step 2: Fitting and selecting appropriate movement models

After range-residency has been confirmed, the next step is to fit a series of range-resident continuous-time movement models to the data, such as the Independent and Identically Distributed (IID), Ornstein-Uhlenbeck (OU; Uhlenbeck & Ornstein, 1930), and OU-Foraging (OUF; Fleming *et al.*, 2014a,b) processes. Model selection should then be employed to identify the model that best fits the data (Fleming *et al.*, 2014b, 2015b, 2019). In `ctmm` this process is carried out via the `ctmm.select()` function.

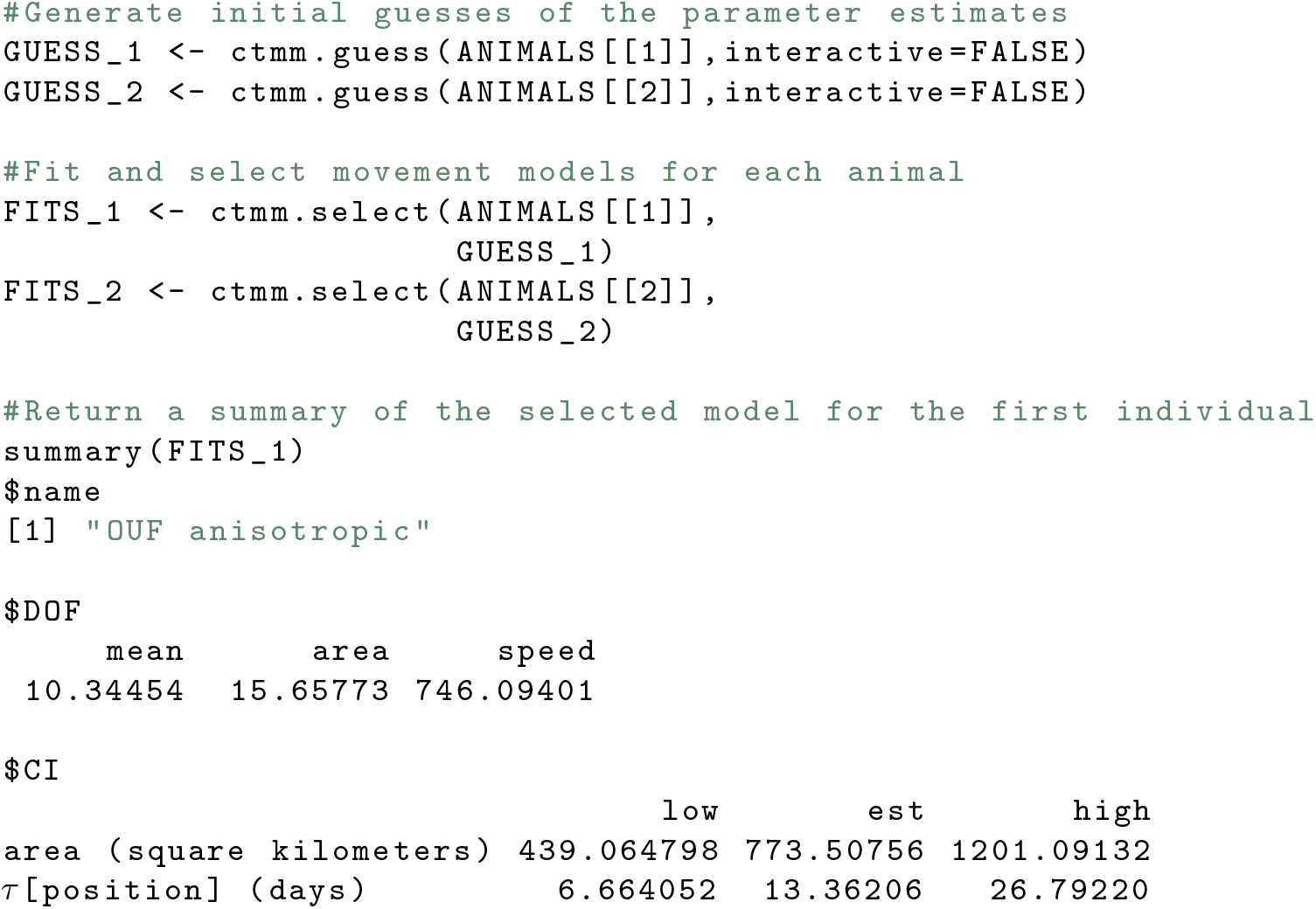
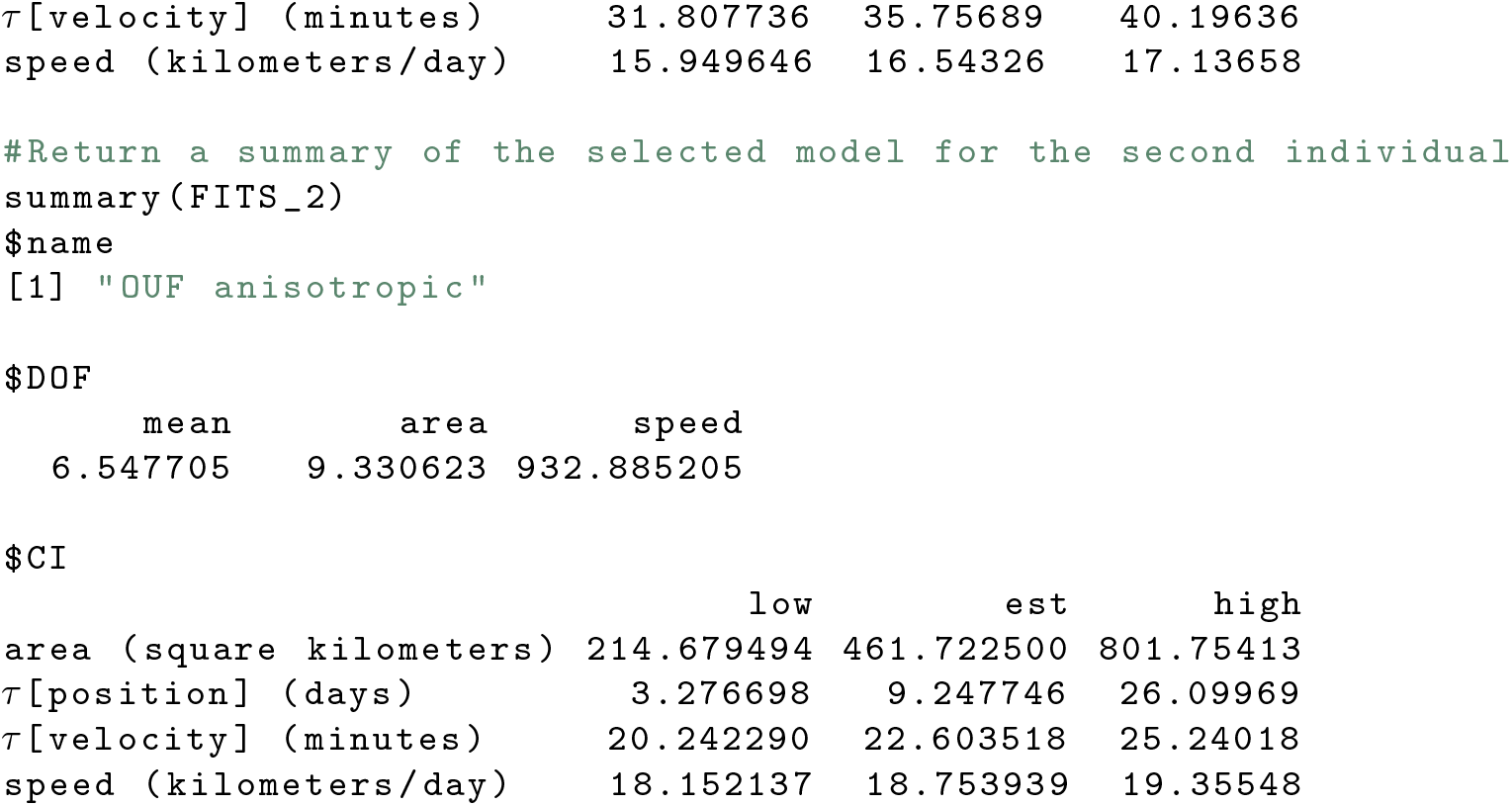

### Step 3: AKDE home range estimation

With a fitted, selected movement model in hand, autocorrelated Kernel Density Estimation (AKDE) home range estimates can then be calculated for the individuals of interest (Fleming *et al.*, 2015a; Fleming & Calabrese, 2017; Fleming *et al.*, 2018). In ctmm this process is carried out via the `akde()` function. Note: It is essential to estimate the HRs for all animals on the same grid so that the CDE can be calculated

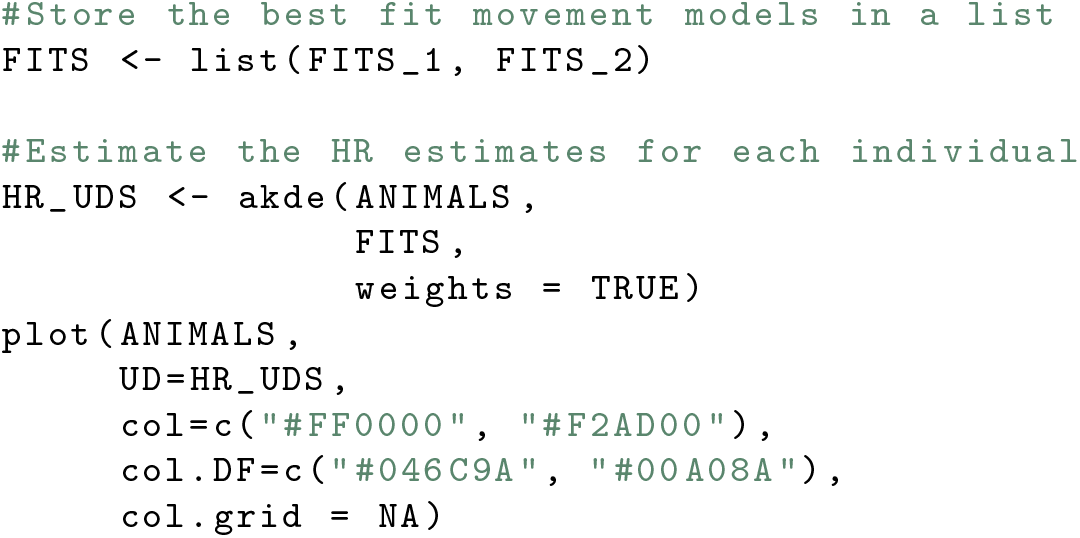

**Figure S5.3:**
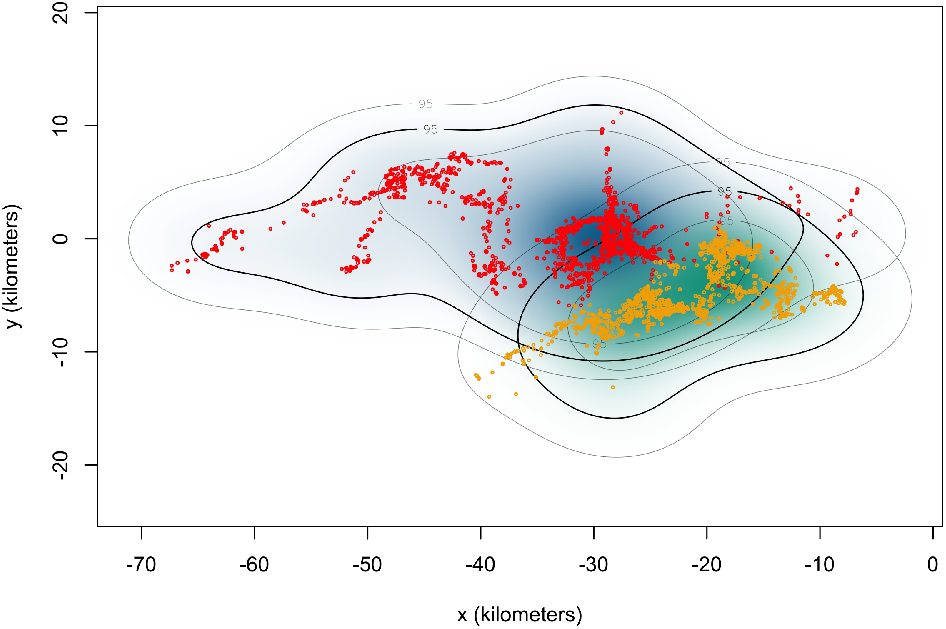
Location data and 95% home range estimates for the two individuals.

### Step 4: CDE estimation

With filtered data, and an appropriate movement model in hand, the final step is to estimate the animal’s instantaneous speeds, the mean speed over the study period, or the speed/distance travelled over a specific period of time.

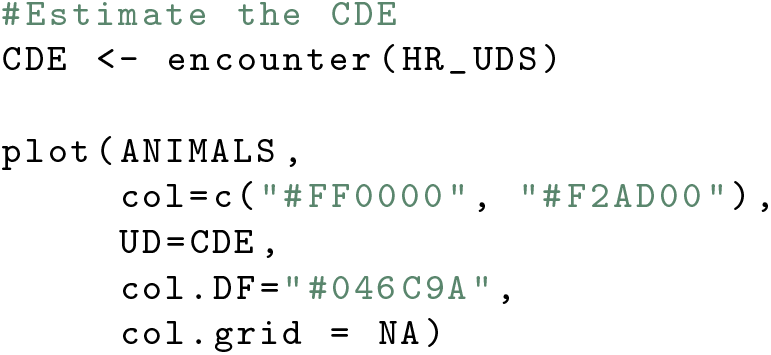

**Figure S5.4:**
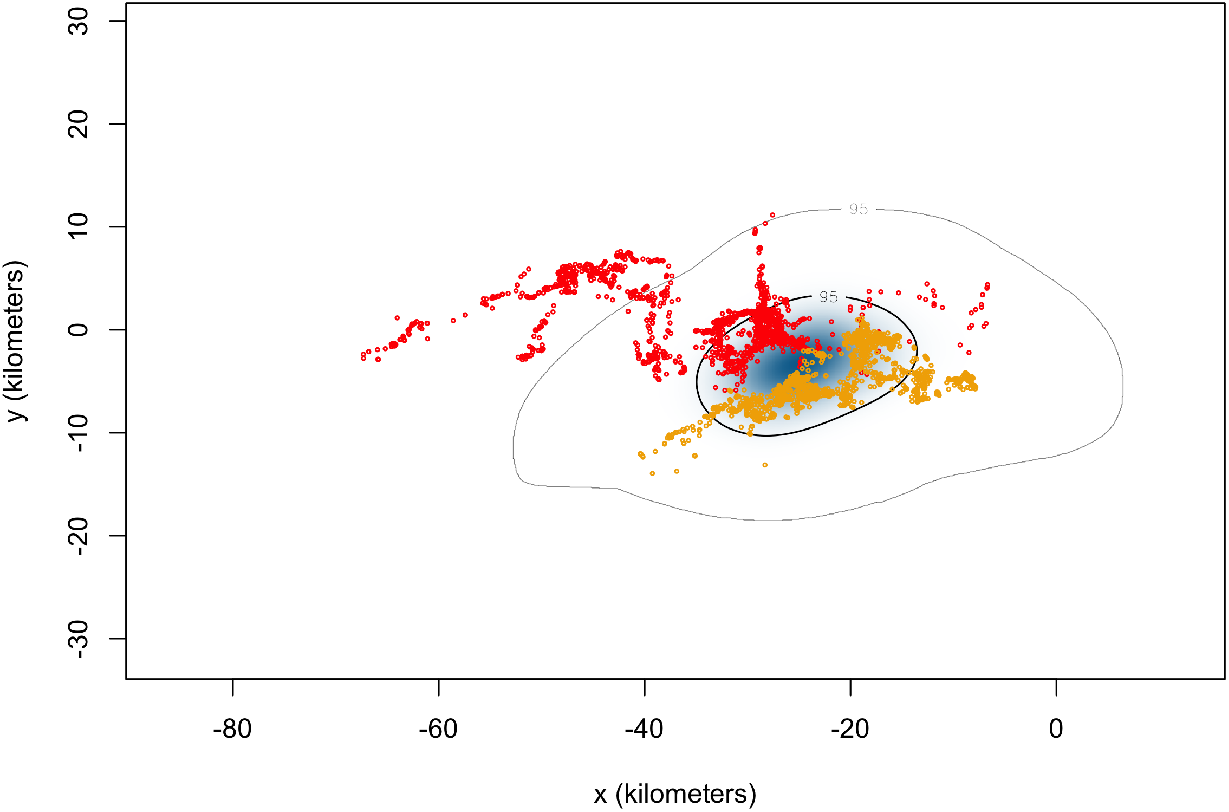
Location data and 95% Conditional Distribution of Encounters (CDE) estimate for the two individuals.

## Footnotes

1 Even though the movement is not isotropic we can still define the *x*-*y* axes in a way that the *y*-axis is aligned with the distance between home-range centers

